# Intuitive and broadly applicable definitions of niche and fitness differences

**DOI:** 10.1101/482703

**Authors:** Jurg W. Spaak, Frederik De Laender

**Affiliations:** University of Namur, Institute of Life-Earth-Environment, Namur Center for Complex Systems, Namur, Rue de Bruxelles 61, Belgium

**Author notes:** Corresponding author, phone-number.

## Abstract

Explaining nature’s biodiversity is a key challenge for science. To persist, populations must be able to grow faster when rare, a feature called negative frequency dependence and quantified as ‘niche differences’ (𝒩) in coexistence theory. Here, we first show that available definitions of 𝒩 differ in how 𝒩 link to species interactions, are difficult to interpret, and often apply to specific community types only. We then present a new definition of 𝒩 that is intuitive and applicable to a broader set of (modelled and empirical) communities than is currently the case, filling a main gap in the literature. Given 𝒩, we also re-define fitness differences (ℱ) and illustrate how 𝒩 and ℱ determine coexistence. Finally, we demonstrate how to apply our definitions to theoretical models and experimental data, and provide ideas on how they can facilitate comparison and synthesis in community ecology.

## Introduction

In order to persist through time, species must exhibit frequency dependence. That is, the population growth of a species must depend on its frequency, i.e. its relative density. Natural communities host a multitude of mechanisms that can lead to frequency dependence. Well-known examples include resource partitioning (Adler *et al*., 2007; Levine & HilleRisLambers, 2009), differential vulnerability to predators (Allan *et al*., 2010; Carson & Root, 2000; Chesson & Kuang, 2008), differential associations with mutualists (Johnson & Bronstein, 2019; Siefert *et al*., 2018) or occupation of distinct microhabitats (Silvertown, 2004). These mechanisms have been collectively coined as ‘niche differences’ (Chesson, 2000; HilleRisLambers *et al*., 2012; Letten *et al*., 2017).

In coexistence theory, one way of quantifying the strength of niche differences is to compare observed population growth with the population growth that is expected when niche differences would be absent (Adler *et al*., 2010, 2007; Chesson, 2000, 2003). Without niche differences, one of the species will eventually exclude all others, where the rate of exclusion depends on the competitive advantage of the winner. This competitive advantage is often called fitness difference (Barabás *et al*., 2018; Chesson, 2000, 2003; Hart *et al*., 2018). A key question is if niche differences in natural systems are sufficiently strong to overcome fitness differences and save species from extinction (Adler *et al*., 2018; Angert *et al*., 2009; Connolly *et al*., 2017; Harris *et al*., 2017; Hubell, 2001; Narwani *et al*., 2013; Usinowicz *et al*., 2017).

Niche and fitness differences formalise species persistence in a way that is conceptual. That is, one does not need to specify the details of the community or its environment, but rather focuses on higher-level processes, i.e. how species grow under different circumstances. This feature would in principle allow synthetic studies across different community types and environmental conditions, with niche and fitness differences acting as common currency that represent the net outcome of detailed ecological mechanisms. Such studies are important because they foster a unified understanding of community composition (Adler *et al*., 2018) and facilitate studying how environmental context and community characteristics jointly influence species persistence, which can help understanding global change effects (Grainger *et al*., 2019).

At present, however, the application of niche and fitness differences is hampered by a lack of consensus on their mathematical definition. Indeed, the operationalisation of these concepts has been discussed for almost a century and new methods are being constantly proposed (Bimler *et al*., 2018; Carroll *et al*., 2011; Chesson, 1990, 2000, 2003; Hurlbert, 1978; Morisita, 1959; Renkonen, 1938), leading to a proliferation of mathematical definitions of niche and fitness differences. We identified 10 definitions available in the literature (appendix A) and found that every single existing definition displays a number of features that limit its applicability. For instance, most of the definitions only apply to communities whose dynamics obey a specific mathematical model (Adler *et al*., 2007; Bimler *et al*., 2018; Chesson, 1990; Chesson & Kuang, 2008; Godoy & Levine, 2014; Saavedra *et al*., 2017). This means that the applicability of these definitions is limited to specific community types. In addition, several definitions cannot be computed for communities with positive species interactions and/or more than two species. Also, not all definitions allow inference of coexistence or exclusion, i.e. niche and fitness differences do not predict whether species will persist or not. Finally, the interpretation of the numbers returned by these definitions has not been examined yet and most likely differs greatly among definitions. This last point is exemplified by the domains of the mathematical definitions (Appendix A) which indeed vary among definitions.

Here, we first show that available definitions of niche differences do not align with biological intuition and present a new definition that does. We also derive the corresponding definition of fitness differences and coexistence conditions. An important feature of these new definitions is that they apply to any mathematical model or empirical system driven by any mechanism, with the sole - but crucial - requirement that invasion analysis correctly predicts coexistence. The flexibility of the new definitions allows comparing different community types, containing an arbitrary number of species and driven by a variety of species interactions, addressing a key limitation in theoretical ecology. Finally, we illustrate theoretical and experimental applications of the new definitions. To this end, we apply the definitions to various models representing a suite of interaction types. We also show how simple growth experiments suffice to quantify niche and fitness differences, using an empirical dataset of two picocyanobacteria competing for light.

## Theory

### A diversity of definitions

To facilitate interpretation and broad application, the definitions for niche and fitness differences should align with biological intuition. That is, intuition dictates that niche differentiation facilitates persistence (𝒩 increases as species persist more easily). Intuition also dictates the values 𝒩 should carry in five specific scenarios. First, when intra- and interspecific interactions are of equal size (*α* = −1 in Fig. 1), individuals of both species are interchangeable: the effect an individual has on another individual does not depend on species identity. Thus, 𝒩 should equal 0 (black triangle in Fig. 1) (Chesson, 1990). Second, when interspecific interactions are absent (*α* = 0 in Fig. 1), each species grows as if other species are absent. Thus, 𝒩 should be some predefined non-zero real number that indicates complete niche differentiation, e.g. 1 (black dot in Fig. 1) (Godoy & Levine, 2014). The third point is the logical consequence of these first two points: intermediate interspecific interaction strengths should result in 𝒩 between 0 and 1 (or some other pre-defined nonzero real number, solid rectangle in Fig. 1). Fourth, when interspecific interactions are stronger than intraspecific interactions, persistence is ‘harder’ (𝒩 should be smaller) than if species occupied exactly the same niche (𝒩 = 0). Consequently, 𝒩 should be negative (dashed rectangle in Fig. 1), as has been stated before (Ke & Letten, 2018; Mordecai, 2011). Fifth, when interspecific interactions are positive, e.g. because of facilitation, the presence of other species makes persistence ‘easier’ (𝒩 should be greater) than if these other species would have no effect on the focal species (i.e. interspecific interactions are absent, in which case 𝒩 = 1). Thus, 𝒩 should inevitably be greater than 1 (dotted rectangle in Fig. 1) when species interactions are positive.

**Figure 1:**
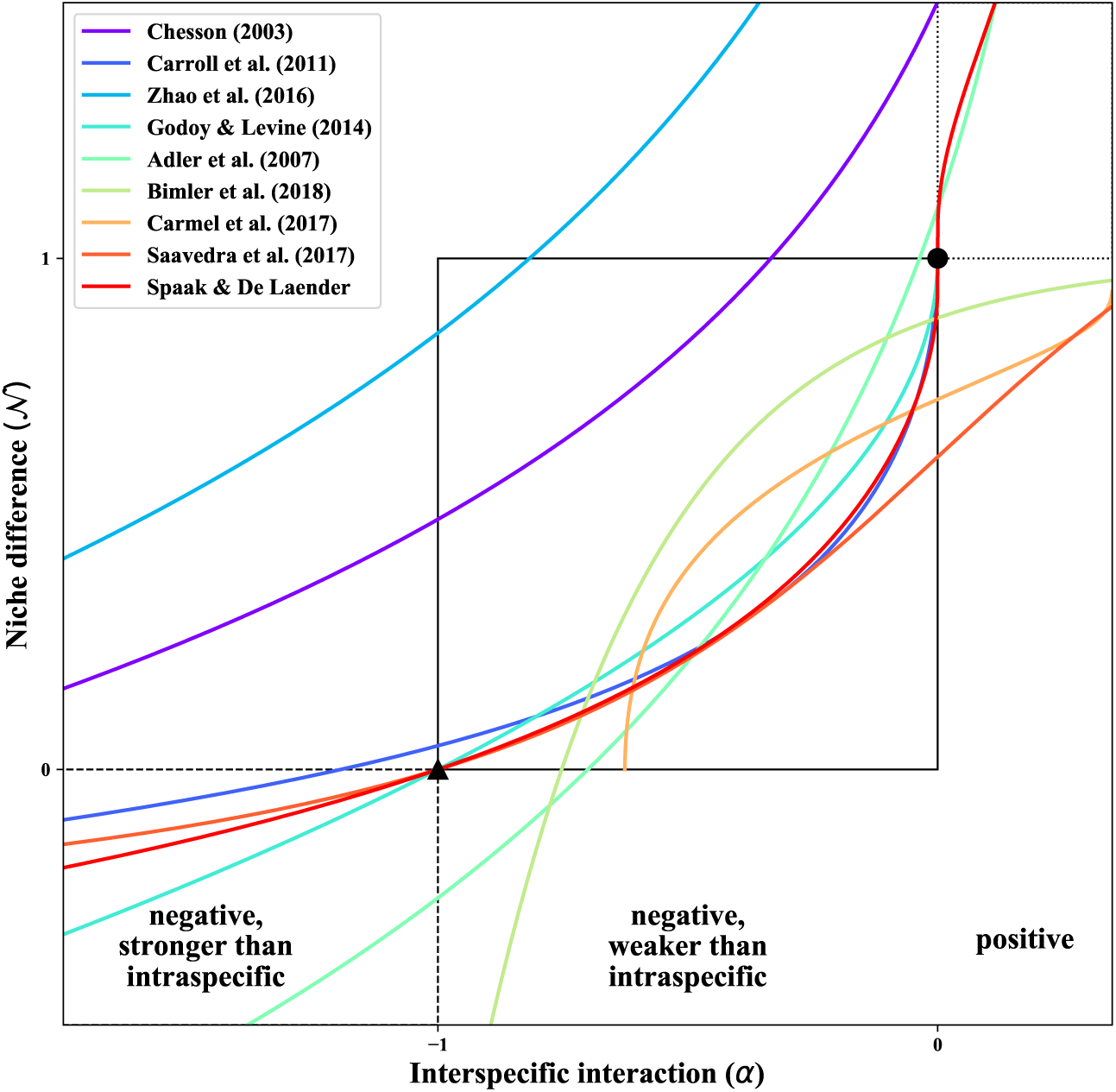
The modelled response of niche differences (𝒩) to the interspecific interaction strength *α* between two annual plants differs among available definitions. The black triangle indicates where inter- and intraspecific interactions are equal (*α* = −1), and so species occupy the same niche, meaning that 𝒩 should be 0. Communities with stronger interspecific interactions must have 𝒩 < 0 (dashed rectangle). The black dot indicates where species do not interact (*α* = 0), and so species have completely different niches, meaning 𝒩 should be 1. Consequently, communities in which interspecific interactions are positive (*α* > 0) should have 𝒩 larger than 1 (dotted rectangle). Finally, for all communities where −1 ≤ *α* ≤ 0, 𝒩 must have intermediate values (0 ≤ 𝒩 ≤ 1, solid rectangle). The new definition proposed here (red), which is applicable to a wide variety of models and experimental data (i.e. not only the annual plant model), complies with this biological intuition. Parameter values, a plot for the corresponding fitness differences (ℱ), and mathematical expressions of the 𝒩 and ℱ definitions are in the appendix A.

We found that available definitions of 𝒩 are unlikely to fulfil the five requirements outlined here. To show this, we computed 𝒩 for the annual plant model, a workhorse of theoretical ecology (Adler *et al*., 2012, 2010, 2007; Angert *et al*., 2009; Germain *et al*., 2016; Godoy *et al*., 2014; Levine & HilleRisLambers, 2009) (Fig. 1), using eight of the ten definitions for niche and fitness differences. The two other definitions cannot be applied to the annual plant model. All definitions return greater 𝒩 as species interactions shift from strongly negative, over weakly negative, to positive. However, different definitions for niche difference imply a variety of niche difference responses to the strength and sign of species interactions (Fig. 1). In addition, these definitions do not map these species interactions to the intuitive niche difference values, as stated above (but see Chesson (1990); Chesson & Kuang (2008); Godoy & Levine (2014)). We therefore introduce, in the next section, a new definition that does align with biological intuition.

### Defining niche differences based on biological intuition

Here, we first construct a general definition for 𝒩 that fulfils the five requirements outlined in the previous section, and is therefore based on biological intuition. To construct a definition of 𝒩, we start by considering the per capita growth of a species *i*

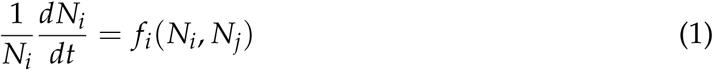

where *N*_*i*_, *N*_*j*_ are the densities of species *i* and species *j* (*i* ≠ *j*) with which *i* interacts. *f*_*i*_ can be essentially any function that describes the per-capita growth rate of species *i*. As done mostly in modern coexistence theory (but see Schreiber *et al*. (2019)), we do not consider Allee effects (positive density dependence), such that we can assume *f*_*i*_(0, 0) > *f*_*i*_(*N*_*i*_, 0): a species grows faster when its density is lower. While this would be technically possible with the definitions proposed here, interpretation of 𝒩 will be challenging (see below). Furthermore, we assume that each species has a stable monoculture equilibrium denoted 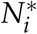 and that the invasion growth rate 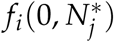 correctly predicts coexistence. That is, the two species *i* and *j* coexist if and only if both species have a positive ‘invasion growth rate’ 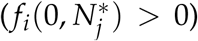. The invasion growth rate is the growth rate of a species when it is reduced to low density (≈ 0) and the other species is at its monoculture equilibrium density. Examples where invasion analysis does not predict coexistence are found in Barabás *et al*. (2018) and Schreiber *et al*. (2019).

When 𝒩 = 0, inter- and intraspecific interactions are equal. Thus, the identity of the individuals does not matter, such that, in eq. 1, *f*_*i*_(*N*_*i*_, *N*_*j*_) is equivalent to writing *f*_*i*_(*N*_*i*_ + *N*_*j*_, 0). However, one cannot simply sum species densities. For example, one tree does not influence nutrient levels as does one forb and will therefore not grow at the same rate. To account for this, we multiply *N*_*j*_ with a factor *c*_*j*_. The ecological interpretation of *c* is discussed below (Applications). Hence, the growth of species *i* can be written as:

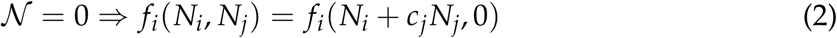

When 𝒩 = 1, interspecific species interactions are absent. Thus, species *j* has no effect on species *i*, and so species *i* grows as if species *j* were absent, i.e. we can put the density of *j* to zero:

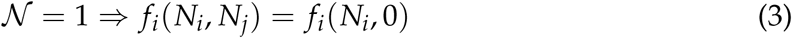

Equations 1-3 hold for all densities *N*_*i*_, *N*_*j*_. However, we will now apply it to obtain species *i*’s invasion growth rate. This corresponds to choosing *N*_*i*_ ≈ 0 and 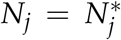, which is *j*’s monoculture equilibrium. In this scenario, eqs. 2 and 3 become 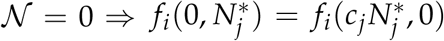 and 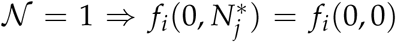. Here, *f*_*i*_(0, 0) is the intrinsic growth rate and 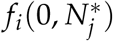 is the invasion growth rate. For 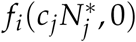, we introduce the term no-niche growth rate of species *i*. This is the growth rate of species *i* if there was no niche differentiation, i.e. if 𝒩 would be 0. The no-niche growth rate of species *i* is the growth rate at the scaled monoculture density of its competitor (species *j*).

The main idea behind the new definitions is to let 𝒩 fulfil the requirements listed in the previous section. The simplest way to do so is by writing 𝒩 as a linear function that equates to 2 and 3 at the desired growth rates:

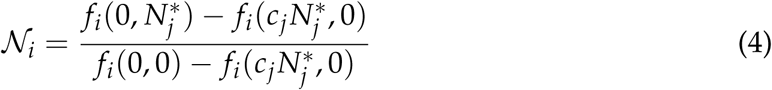

This new definition by design fulfils the requirements listed before, which can be seen when applying it to the annual plant model as done for the existing definitions (Fig. 1). When species interact negatively and do so more within than between species, 𝒩_*i*_ is bounded in [0, 1] (solid rectangle). When interspecific interactions are stronger than intraspecific interaction 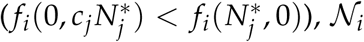 will be negative (dashed rectangle). When interspecific effects are positive 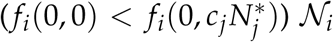 is larger than 1 (dotted rectangle).

This new definition should be interpreted as follows. The numerator of 𝒩_*i*_ compares the growth of species *i* when only interspecific interactions are present 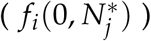 with its growth when only intraspecific interactions matter 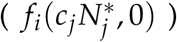. Note that in this last growth rate, 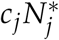 denotes a density of species *i*. Both growth rates are evaluated at the same total scaled density, but at different frequencies of species *i*, being 0% in 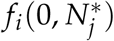 and 100% in 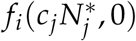. The numerator of 𝒩_*i*_ therefore effectively measures frequency dependence of species *i*’s (Adler *et al*., 2007; Levine & HilleRisLambers, 2009). The denominator of 𝒩_*i*_, which is always positive and thus does not influence the sign of 𝒩_*i*_, compares the growth of species *i* when its density is ≈ 0 with its growth when its density is at the scaled equilibrium density of 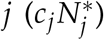. Thus, the denominator of 𝒩_*i*_ measures the strength of species *i*’s density dependence. 𝒩_*i*_ therefore measures the strength of frequency dependence, relative to that of density dependence. According to this new definition, and unlike almost all other definitions (but see Adler *et al*. (2007)), 𝒩_*i*_ is species-specific and is therefore not a community characteristics. However, 𝒩_*i*_ does depend on species *j* as well, as species *j* will influence species *i*’s invasion and no-niche growth rates (eq. 4). In what follows, we use the subscript *i* (𝒩_*i*_) only to distinguish between the niche differences of the species, and use 𝒩 to refer to niche differences in general.

### Fitness differences and coexistence

The novel definition of 𝒩 implies a new definition of the fitness difference ℱ. Verbally, ℱ should represent the per-capita growth rate when both species occupy the same niche, i.e. when 𝒩 = 0 (Adler *et al*., 2010; Barabás *et al*., 2018). Therefore

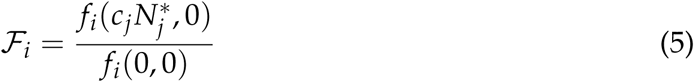

ℱ_*i*_ ranges from −∞ to 1 (because we assume no Allee effects, i.e. 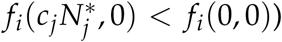 and measures how well species *i* grows in the absence of frequency dependence (no-niche growth rate, numerator) (Adler *et al*., 2010, 2007), compared to its intrinsic growth rate (denominator). When ℱ_*i*_ is 0, species *i* is equally competitive as species *j*. Otherwise exactly one species, the competitive dominant, has ℱ_*i*_ > 0.

Now that we have defined both 𝒩 and ℱ, we can evaluate when species *i* can coexist with species *j*. Interestingly, normalising the invasion growth rate by the intrinsic growth rate yields 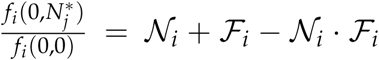 (Appendix B). Thus, *i* can persist within the community when^1^:

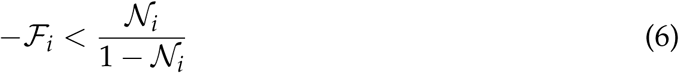

This inequality formalizes the idea that species persist, when 𝒩 “overcome” ℱ. However, the inequality is only meaningful if invasion growth rate correctly predicts coexistence. This inequality yields a number of important insights. First, as for 𝒩, also ℱ is species-specific. Taken together, this shows that the above inequality should therefore be considered as the condition for species *i* to persist. Only if all species from a community fulfil this inequality, species will coexist. Second, the minus sign on the left hand side shows that a high ℱ_*i*_ implies a competitive advantage for species *i*, which is consistent with previous insights (Adler *et al*., 2007; Chesson, 2000, 2003). Third, completely different niches are sufficient to overcome arbitrarily large ℱ_*i*_ (i.e. 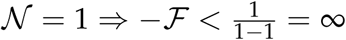). Conversely, if species occupy the same niche (i.e. 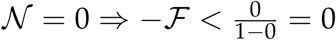), coexistence is only possible under neutrality (i.e. ℱ_*i*_ = ℱ_*j*_ = 0). Fourth, species with negative 𝒩 cannot coexist, as species’ growth is positively frequency dependent: species grow faster when abundant (Ke & Letten, 2018; Mordecai, 2011; Schreiber *et al*., 2019).

### Extension beyond species pairs

The definitions for 𝒩 and ℱ naturally extend to communities composed of more than two species, hereafter ‘multispecies communities’. To show this, we generalised the invasion growth rate and the no-niche growth rate to the case of multispecies communities (for technical details, see Appendix B):

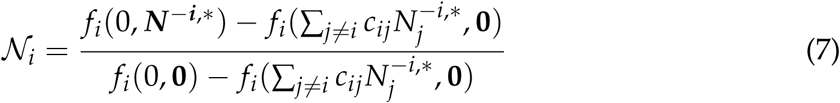

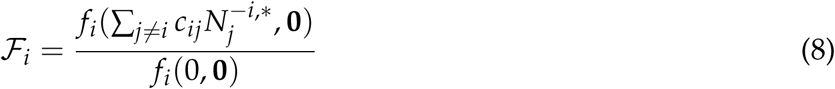

Here ***N***^−*i*,*^ is the vector of equilibrium densities in the absence of species *i*, **0** denotes the absence of all species other than *i*, and similar to the definition for species pairs (eq. 4), *c*_*ij*_ converts densities of species *j* into *i*,. These definitions measure the net effect of species interactions on 𝒩 and ℱ, i.e. direct, indirect (Godoy *et al*., 2017) and higher order effects (Grilli *et al*., 2017). Importantly, the interpretations given for the two-species community still apply, given that (i) invasion analysis is possible and (ii) correctly predicts coexistence (Chesson, 1994, 2000; Turelli, 1978). We acknowledge that, in multispecies communities, (i) and (ii) are sometimes not met (Barabás *et al*., 2018; Saavedra *et al*., 2017).

## Applications

### Application to community models

A first step in applying eqs. 4 and 5 to a model is the quantification of the factors *c*_*i*_ and *c*_*j*_. The *c* convert species *i* to *j* and vice-versa, and so logically *c*_*j*_ · *c*_*i*_ = 1. For example, if one tree influences resource levels ten times more than a forb (*c*_*tree*_ = 10), the forb influences resource levels ten times less than the tree (*c* _*f orb*_ = 1/10). After scaling, both species thus have the same total influence on the environment. In Fig. 2A, we provide an example of two species consuming common resources. We scaled their consumption rates such that total consumption is the same for both species: the white and the grey area are equal. This example shows that both species now also happen to have the same proportion of shared limiting factors (1 − 𝒩_*i*_ = light grey region = 1 − 𝒩_*j*_). We can therefore find *c* by solving the equations

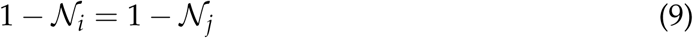

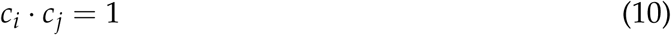

**Figure 2:**
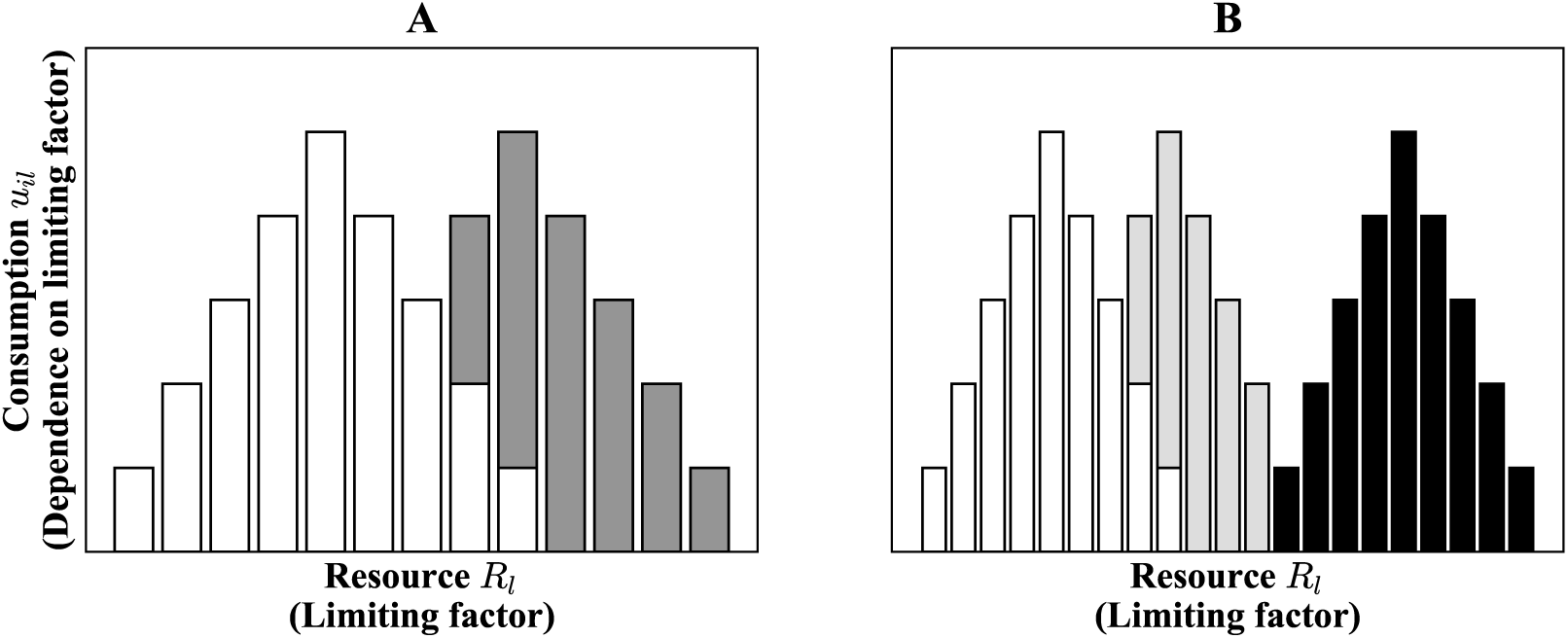
Influence on limiting factors (here, resources) for a two (A) and a three (B) species community. In the two-species community (A) both species have the same total influence on the limiting factors (i.e. *c*_*i*_ = *c*_*j*_ = 1), therefore the amount of shared resources (1 − 𝒩_*i*_ = light grey area = 1 − 𝒩_*j*_) is identical for both species. This is, however, not the case in a multispecies community (B) (Adler *et al*., 2007), where the amount of shared resources is smaller for the black species than for the white species, even though they both consume the same total amount of resources. We therefore expect 𝒩_*black*_ ≠ 𝒩_*white*_.

In Box 1, we illustrate this first step, and the calculation of 𝒩 and ℱ, for a MacArthur consumer-resource model. We then convert this model into the well-known Lotka-Volterra model to express 𝒩 and ℱ using interaction coefficients. This exercise highlight the following results. First, while 𝒩 and ℱ are species-specific, they can be identical between species in species pairs competing for shared resources. Indeed, changing *i* for *j* in eq. 18 shows that 𝒩_*i*_ = 𝒩_*j*_. However, they cease to be identical when including more than two species, as can be seen from Fig. 2B. Indeed, niche overlap, and therefore 𝒩, is species-specific in that case. Second, the new definitions of 𝒩 and ℱ, when applied to the Lotka-Volterra model, collapse to the same definitions for 𝒩 and ℱ previously found for the same model (Chesson, 1990). This shows that these new definitions, which apply to any model (for which invasion analysis is possible and useful) still agree with the definitions found for this particular model. Third, *c*_*i*_ carries a biological interpretation: in the MacArthur model, *c*_*i*_ indeed scales with the total influence on limiting factors.

#### Box 1

𝒩 and ℱ for the MacArthur and Lotka-Volterra model

Consider a community of two species whose dynamics follow (MacArthur, 1970)

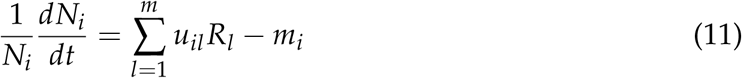

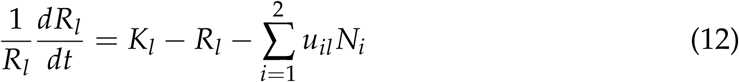

Where *u*_*il*_ is the rate at which species *i* consumes resource *l, R*_*l*_ is the density of resource *l, m*_*i*_ is the loss rate and *K*_*l*_ is the resource’s carrying capacity. We assume that the resource dynamics are faster than the dynamics of the consumers, such that *R*_*l*_ is always at equilibrium. In that case, the model simplifies to (MacArthur, 1970):

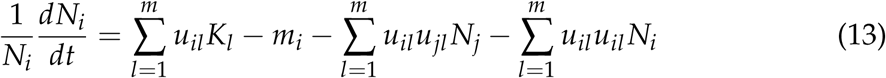

Solving equations 9 and 10 yields 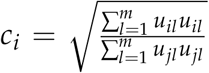 (appendix C). Thus, *c* indeed scales the species’ total influence on limiting factors. Replacing *c*’s into the growth rates, one obtains (Appendix C):

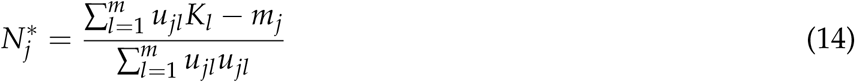

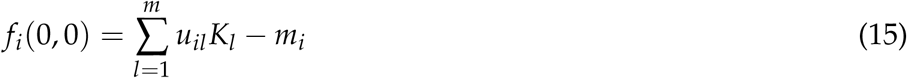

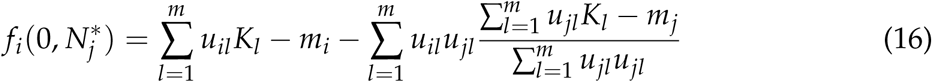

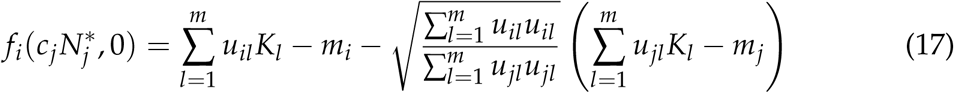

Finally, replacing these into eqs. 4 and 5, one obtains (Appendix C):

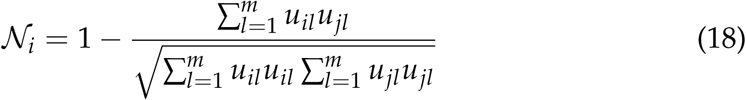

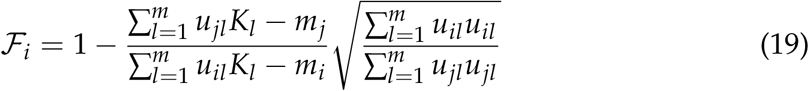

We now note that eq. 13 is equivalent to the Lotka-Volterra model 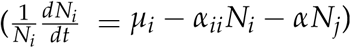, where 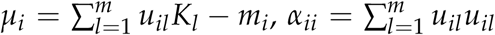, and 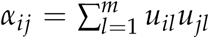 are the intrinsic growth rate, the intraspecific interaction strength, and interspecific interaction strength, respectively. Plugging these expressions in eqs. 4 and 5 recovers the well known equations for 𝒩 and ℱ in the Lotka-Volterra model Chesson (1990, 2000, 2013):

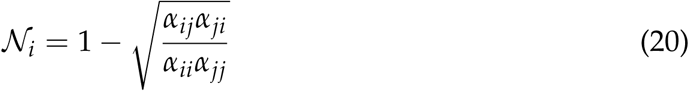

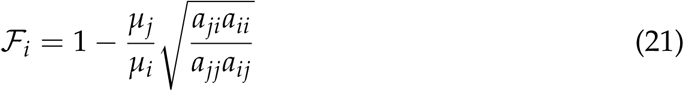

This last feature is independent of the specific model formulation, i.e. it extends beyond the McArthur resource model to any model in which two species interact through resource consumption, resource consumption stimulates growth, and species consume higher amounts of resource when resource availability is higher. In appendix C, we show a mathematical proof that in such a model, increasing the resource consumption of species *i* will increase *c*_*i*_, i.e. *c* is linked to the total resource consumption of a species. Finding the *c* when species have positive effects on each other (for example by generating resources or by limiting the efficacy of a predator) requires additional considerations, which are discussed in appendix B and C.

Finally, we apply equations 4 and 5 to examine how the various growth rates underlying 𝒩 and ℱ, as well as 𝒩 and ℱ itself, change across community types (Fig. 3) modelled using Lotka-Volterra equations (Appendix C). Priority effects occur when interspecific interactions are stronger than intraspecific interactions, i.e. 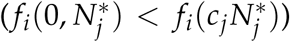. Neutrality occurs when 𝒩 = ℱ = 0 (Adler *et al*., 2007). Competitive exclusion represents the well-known case where 𝒩 are not large enough to compensate for ℱ: only the competitive dominant persists (Chesson, 2013; Ke & Letten, 2018). For the case of parasitism and mutualism, one or both species have an invasion growth rate that is higher than their intrinsic growth rate, respectively: these species profit from other species and thus grow better together than alone. Therefore, these species have 𝒩 > 1. In these cases, ℱ matter less for persistence (they only indicate the winner when 𝒩 = 0) because the coexistence region increases rapidly with 𝒩.

**Figure 3:**
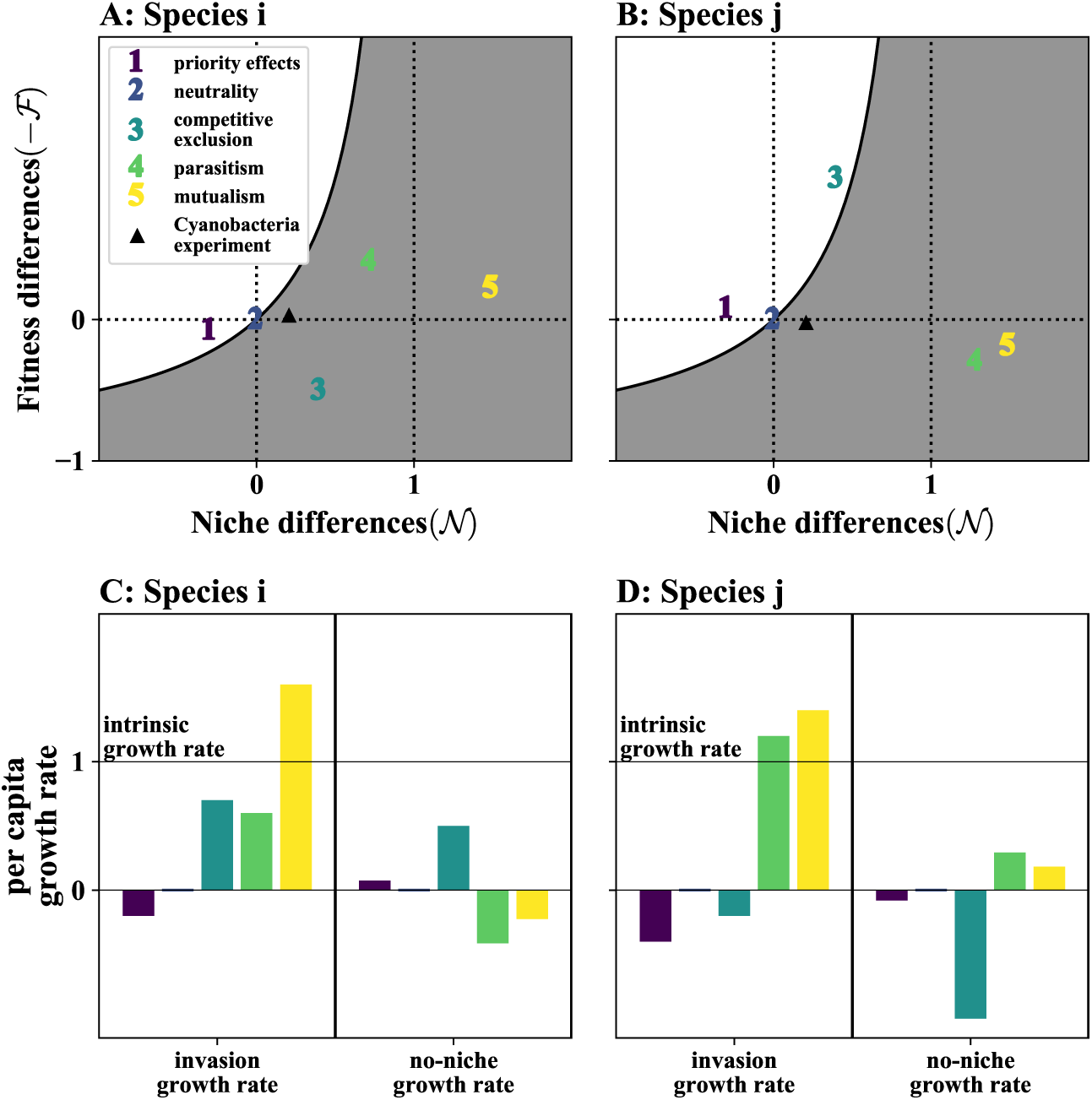
Example computation of 𝒩 and ℱ for common two-species communities. A and B show the distribution of 𝒩 and ℱ for species i and species j respectively, where color codes refer to different communities (see legend). 1-5 are communities simulated with Lotka-Volterra models, while ‘experiment’ refers to the performed experiment (Fig. 4). Species in the grey area have a positive invasion growth rate, i.e. they persist. If both species have positive invasion growth rates the species are assumed to coexist (Barabás *et al*., 2018; Chesson, 2000). C and D compares the invasion and the no-niche growth rate to the intrinsic growth rate (shown by the vertical full line). This comparison gives qualitative insight (e.g. the sign) on 𝒩 and ℱ.

### Application to experiments

The applicability of the new 𝒩 and ℱ definitions extends beyond models and can be used to analyse coexistence empirically. In these experiments, one needs to measure the various growth rates present in equations 4 and 5 to quantify 𝒩 and ℱ (Fig. 4). These experiments also allow estimating the factors *c*_*i*_ and *c*_*j*_, giving insight in the species’ total influence on limiting factors. Importantly, the definitions can be computed directly from the measured growth rates, without any assumption on the species’ ecology or the need to fit a model, contrary to many other definitions 𝒩 and ℱ. This is particularly useful since natural communities are typically governed by a multitude of species interactions, many of which will be unknown (Carrara *et al*., 2015; Montoya *et al*., 2006).

**Figure 4:**
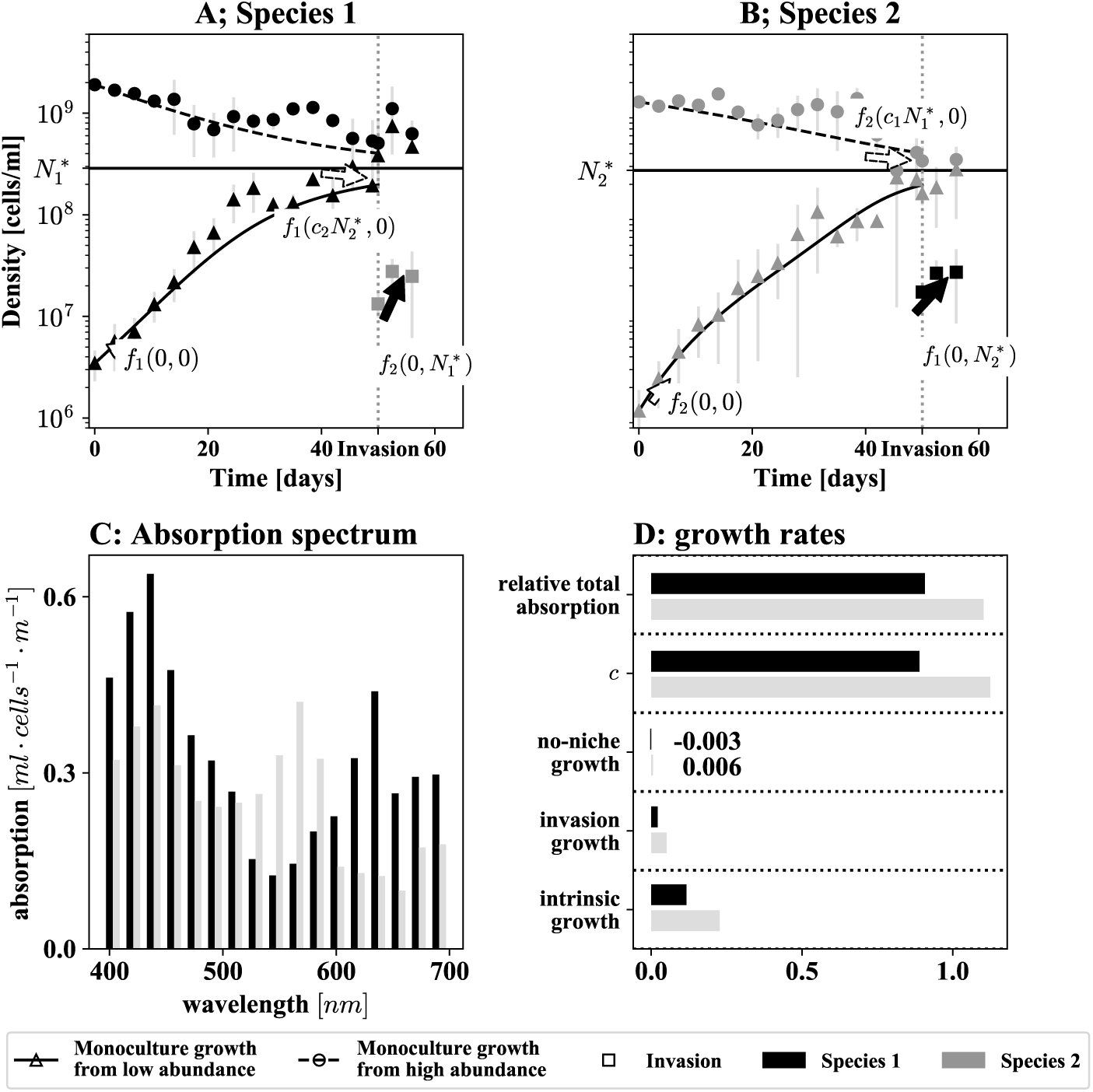
We measured 𝒩 and ℱ for two marine cyanobacteria species from the genus *Synechococcus*, sampled in the Baltic sea (Stomp *et al*., 2004). A and B: Population growth in the different experiments with different starting conditions. The arrows indicate the growth rates we measured to quantify 𝒩 and ℱ. Error bars (grey) show one standard deviation (3 replicates). C: The two species have different absorption spectra and therefore partition light usage. D: The experiment confirms that the species compete and coexist, as the invasion growth rate is positive, but smaller than the intrinsic growth rate. The conversion factor *c* is very similar to the relative total absorption of the two species, confirming the theory. An automated code to compute 𝒩 and ℱ from such experimental data can be found on https://github.com/juergspaak/NFD definitions.

To illustrate the application to experimental data, we performed an experiment in which we measured growth of two picocyanobacteria species competing for light (Fig. 4). Detailed experimental methods can be found in the appendix D. The two picocyanobacteria species contain different pigments (phycocyanobilin and phycoerithrobilin), which allow them to absorb different wavelengths of light (Fig. 4 C). Because light colour usages of these two species partly overlap, exactly as did resource usage in the MacArthur model (Fig. 2), we expected that 0 < 𝒩 < 1 (i.e. species compete). Experiments and field data have shown that pigmentation differences among picocyanobacteria lead to a resource (light) partitioning that is sufficiently strong to allow coexistence (Stomp *et al*., 2004, 2007a,b). We therefore also expected that 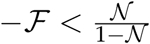 (i.e. coexistence).

Three growth curves per species suffice to quantify 𝒩 and ℱ for a two-species community (Fig. 4). First (Fig. 4A and B, triangles), we grew both species in a monoculture, starting from low density to obtain the intrinsic growth rate. Second (Fig. 4A and B, circles), we grew both species in a monoculture starting from a density higher than their equilibrium density to obtain the no-niche growth rate. In this experiment, the growth rate at which the density of the focal species reaches that of the scaled equilibrium density of its competitor 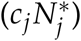, is the no-niche growth rate. Third (Fig. 4A and B, squares), we introduced each of both species into a monoculture at equilibrium of its competitor to obtain the invasion growth rates. More precisely, we introduced 5% of the invading species’ equilibrium density (Gallego *et al*., 2019; Narwani *et al*., 2013). We estimated all these growth rates as 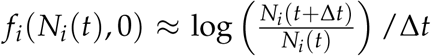 with Δ*t* = 84 hours. We then fitted a univariate spline to estimate these growth rates at the various densities. Finally, we were able to use all these growth rates to solve the equation 9 and thus obtain *c*_*i*_ and *c*_*j*_, as well as 𝒩 and ℱ. Importantly, the scaled equilibrium density at which the no-niche growth rate is measured is part of the solution to these equations.

The results of the experiment confirmed our expectations: species compete for light (0 < 𝒩 < 1 for both species) and coexist (see triangle in Fig. 3). The estimated growth rates show that both species can grow independently of each other (positive intrinsic growth rate), and can invade each other’s monoculture (positive invasion growth rate). Their no-niche growth rate is much smaller than their corresponding intrinsic growth rates, and slightly negative for species 1 but positive for species 2. This shows that removing all niche differentiation would lead to the exclusion of species 1, as is also seen from these species’ fitness differences ℱ (Fig. 3). Finally, we found the conversion factors *c*_*i*_ and *c*_*j*_ to match the relative total resource consumption (absorption) of the two species (figure 4 D). This finding aligns with the theoretical result that the conversion factors link to the total influence on limiting factors (available resources) and confirms that these species compete for light.

While this experimental procedure is applied to fast growing communities, this design can be applied to communities with slow growing species as well. Any method that allows estimating per-capita growth is sufficient, but obviously these methods will vary with the considered community. E.g. for annual plants, one may sow different quantities of seeds, ranging from low to above equilibrium density, in plots, and measure their growth.

## Discussion

In this article, we propose new definitions for 𝒩 and ℱ that are biologically intuitive by design. The approach is similar to Carroll *et al*. (2011) in that it allows computing 𝒩 and ℱ from simulations or experimental data, without the knowledge of the underlying mechanisms. When applied to the Lotka-Volterra model for competing species, the definitions collapse to the same mathematical expressions of 𝒩 and ℱ found before (Chesson, 1990, 2013), while still being applicable to a large body of community models. This indicates that there is a potential for these new definitions to unify existing definitions (Barabás *et al*., 2018; Carroll *et al*., 2011; Chesson, 2000; Godoy & Levine, 2014), while enforcing the connection between theory and biological intuition (Adler *et al*., 2010, 2007; HilleRisLambers *et al*., 2012).

### Specificities and limitations

𝒩 and ℱ, as defined in this paper, differ from other definitions of niche and fitness differences. Most notably, the proposed definitions are not based on specific mathematical models, apply to communities with positive species interactions and/or more than two species, and allow inference of coexistence or exclusion. Thus, the new definitions notably extend coexistence theory based on invasion analyses. The structural approach of Saavedra *et al*. (2017) is the only definition for niche and fitness differences which can analyse communities that are outside of the scope of this new definition, as it does not depend on invasion analysis. They define 𝒩 and ℱ for any community in which the equilibrium point of the community can be described as ***r*** = ***αN***\*, where ***α*** is a *n* by *n* matrix containing the species interactions and ***r*** is a vector containing the intrinsic growth rates (or equivalent). Finally, there are still communities that are beyond the reach of all definitions for 𝒩 and ℱ, including the newly proposed definitions: multispecies communities with non-linear species interactions (therefore excluding the approach of Saavedra *et al*. (2017)) and not allowing invasion analysis (therefore excluding the approaches of Carmel *et al*. (2017); Carroll *et al*. (2011); Chesson (2003) and the definitions proposed here).

The reliance on invasion analysis is a first limitation of the proposed definitions, as it is for many other definitions (Carmel *et al*., 2017; Carroll *et al*., 2011; Chesson, 2003; Zhao *et al*., 2016). This reliance means that one should be able to compute the invasion growth rate for each species and that the invasion growth rates correctly predict coexistence. This can limit the applicability of the definitions in two ways. First, there will be communities in which invasion analysis does not correctly predict coexistence (Barabás *et al*., 2018). An example is the annual plant model combined with positive frequency dependency proposed by Schreiber *et al*. (2019). Species in this community never have a positive invasion growth rate, but there may be a feasible and stable two-species equilibrium point (Schreiber *et al*., 2019). Second, invasion analysis requires that all species within each S-1 subcommunity (the community without the invading species) stably co-exist. A well-known counter example is the rock-paper-scissors community, in which the whole community can coexist, while each two-species subcommunity is not stable (Grilli *et al*., 2017). While these two assumptions will be met for most two-species communities, we expect they will be increasingly violated as communities contain more species (Saavedra *et al*., 2017).

A second limitation of the new definitions is the difficulty of interpretation that arises in communities with Allee effects. The proof that the *c*_*i*_ have a unique solution demands Allee effects to be absent (see Appendix B). Consequently, Allee effects imply that species may have multiple 𝒩 and ℱ. This highlights the meaning of Allee effects: species change their dependence on limiting factors with their density. While the new definitions do allow computing these multiple 𝒩 and ℱ, it is at present not clear how to interpret them.

### The need for new definitions

With already ten definitions at hand, one may ask why we need new definitions for niche and fitness differences. We identify at least two reasons. A first reason deals with the complexity of many community models. Many approaches to compute niche and fitness differences first fit a community model to empirical data and then perform maths to link the model to 𝒩 and ℱ (Bimler *et al*., 2018; Chesson, 1990; Godoy & Levine, 2014; Saavedra *et al*., 2017). One challenge is that these maths are often non-trivial (e.g. Carmel *et al*. (2017); Godoy & Levine (2014); Saavedra *et al*. (2017)) and one needs to resort into simplifying the community model (Godoy & Levine, 2014; Letten *et al*., 2017). This may lead to the omission of mechanisms contributing to 𝒩 (Chu & Adler, 2015). For example, niche partitioning could arise at different life stages of a species (Moll & Brown, 2008), or through its interactions with resources (Chesson, 1990), predators (Chesson & Kuang, 2008) or mutualists (Johnson & Bronstein, 2019) and will be affected by environmental change (Rey *et al*., 2017; Wainwright *et al*., 2018). An important advantage of the definitions is that they do not require analytical solutions of a community model or even a community model at all: one can simply simulate or perform the experiments described in the section “Application to experiments” and measure the resulting growth rates to compute 𝒩 and ℱ. Thus, the model or experimental community can be used in its full complexity, capturing all mechanisms potentially contributing to 𝒩 and ℱ.

A second reason is that the analysis of communities with non-competitive interactions (e.g. mutualistic and trophic, Fig. 1) and multiple species (eq. 7) is urgently needed. Indeed, such communities have often been analysed in a suboptimal way. For example Narwani *et al*. (2017) tested whether closely related fresh water green algae are more likely to coexist due to higher niche differentiation. However, 𝒩 could not be computed when species interactions were positive. Similarly, in a meta-analysis on terrestrial plants, Adler *et al*. (2018) were not able to compute 𝒩 for one third of the data, as they contained positive interactions. Chu & Adler (2015) measured 𝒩 and ℱ in an age structured model for perennial plants fitted to long-term demographic data, Petry *et al*. (2018) measured the effects of ant consumption on 𝒩 and ℱ and Veresoglou *et al*. (2018) reanalysed data from the “BIODEPTH” grassland biodiversity experiment. While these studies do report computed 𝒩 and ℱ for multispecies communities, the interpretation of these variables is difficult, as they do not predict coexistence in multispecies communities.

### New insights and outstanding questions

Historically, 𝒩 measured the proportion of resources not shared by two species (Hurlbert, 1978). Being a proportion, 𝒩 was bound between 0 and 1 (Godoy & Levine, 2014). Linking a mechanistic (resource uptake) model to the Lotka-Volterra model (Chesson, 1990; MacArthur, 1970) was a key step in exploring 𝒩 beyond the traditional range [0, 1]. Recent research interpreted negative 𝒩 as a sign that interspecific interactions are stronger than intraspecific interactions, leading to priority effects (Grainger *et al*., 2019; Ke & Letten, 2018). The interpretation that 𝒩 greater than 1 imply positive interspecific interactions is a logical next step. Our results show that this interpretation is correct when both species have symmetric positive effects on each other, but also that species benefiting from other species (e.g. parasitism in Fig. 3) would have 𝒩 > 1.

The results suggest that 𝒩 and ℱ are species-specific properties. While this idea has already been introduced by Adler *et al*. (2007), virtually all other definitions consider 𝒩 a community property. This likely stems from the fact that most definitions focus on two species communities with competitive interactions, in which case niche differences are the proportion of shared resources, which is the same for both species (see Fig. 2 A, light grey area). Therefore, in this particular case, the two species have the same 𝒩, leading to the impression that 𝒩 is a community property.

The results spur three outstanding questions on species coexistence. A first question deals with the variable *c*, that we found scales with the total influence on limiting factors, both for a class of resource competition models and empirically. However, our mechanistic understanding of these factors is absent for models beyond the ones considered here, notably in systems not driven by resource competition. Most notably, we do not know how the *c* relate to the presence of limiting factors that have negative effects on per-capita growth. A second outstanding question deals with the location of species from complex communities on the 𝒩 and ℱ plane from Fig. 3. While these positions may be trivial in some two-species communities, they will not be in large complex networks with a high number of indirect effects, possibly leading to surprising conclusions regarding the contribution of stabilizing and equalizing forces to persistence. A third question deals with the extended applicability the new definitions offer to coexistence theory (as long as invasion analysis is possible and useful). This applicability would allow asking how 𝒩 or ℱ compare across community types, mechanisms, and environments. Thus, the new definitions enable cross-community comparisons in a way that at present is not possible. One could, for example, examine how species from different community types position in Fig.3, to ask if community types that are thought to harbour a more diverse set of mechanisms fostering coexistence (e.g. annual plants) distinguish from community types that appear to have little possibilities for niche differentiation (e.g. phytoplankton (Hutchinson, 1959)).

Within a community type (e.g. phytoplankton), one could compare the stabilizing effect of various mechanisms. For example, we found 𝒩 and ℱ to indicate coexistence in a classic example of a community driven by partitioning of the light spectrum through phenotypic differences (i.e. pigmentation, see Fig. 3) (Stomp *et al*., 2004). How does the stabilizing strength of these phenotypic differences (driving 𝒩) compare to the strength of other relevant mechanisms (e.g. competition for mineral nutrients, allelopathy)? One could also examine how environmental changes that alter the sign of species interactions (Olsen *et al*., 2016) impact the persistence, since the proposed definitions accommodate various interaction types.

In conclusion, our results offer a new perspective on two concepts that underpin biodiversity science, and foster an intuitive biological interpretation of how similarities and differences among species map to the persistence of species (Fig. 1). The developed theory is applicable to a variety of ecological communities, regardless of community complexity, and without the need of mathematical skills (Ellner *et al*., 2018), for any system in which invasion analysis is possible and correctly determines coexistence. The fact that all these communities can be analysed with one approach is a major step forward. Taken together, the novel definitions of 𝒩 and ℱ we present here promote conceptual unification and facilitate empirical research in community ecology and biodiversity science.

## Supplementary Information

An automated code that will compute 𝒩 and ℱ for any given ecological model or experimental data is available. The code is available in Python and in R on https://github.com/juergspaak/N

## Acknowledgments

We thank O. Godoy, G. Barabas and S. Ellner for comments on earlier versions of this manuscript. We thank J. Virgo for conducting the experiment. F.D.L. received support from grants of the University of Namur (FSR Impulsionnel 48454E1), and the Fund for Scientific Research, FNRS (PDR T.0048.16).

## Footnotes

1 Assuming that 𝒩_*i*_ < 1

