## Appendix for "Intuitive and broadly applicable definitions of niche and fitness differences"

### Appendix A: Review

As mentioned in the main text the different available definitions all suffer from one or several drawbacks which we want to highlight here in the appendix. Table A1 summarises many features of the different definitions.

The mathematical equations for each definition are given in the column *Definition*. A subscript $i$ indicates that the parameter is species specific. Note that we log-transformed the definition of Adler *et al.* (2007) to be able to compare the different definitions more consistently. Furthermore for the Saavedra *et al.* (2017) method only the definition for two-species community is given, for the multispecies case we refer to their original paper.

The *Range* column explains the range of the definitions for the communities without facilitation and with intraspecific competition being stronger than interspecific (solid rectangle in fig. 1 from the main text). In the Lotka-Volterra and annual-plant setting this means $\alpha_{ij}\alpha_{ji} \leq \alpha_{ii}\alpha_{jj}$. As argued in the main text, this range should be [0,1].

The *Pos. Eff.* contains the range of $\mathcal{N}$ when species do exhibit positive interactions. *Undef* indicates that $\mathcal{N}$ and or $\mathcal{F}$ are not defined for this case, e.g. because one would have to take the square root of a negative number. Numbers in red indicate that the range overlaps with the usual range. The definition of Saavedra *et al.* (2017) is defined for small positive interspecific interactions (for which the range is given), not however for large. The *Comp.* contains the range of $\mathcal{N}$ when interspecific interactions are stronger than intraspecific interactions.

Given the $\mathcal{N}$ and $\mathcal{F}$ values the *Coex.* indicates what relation must be fulfilled in order to have coexistence. For some definitions the knowledge of $\mathcal{N}$ and $\mathcal{F}$ however is not sufficient to infer coexistence.

The column *Add.* indicates some additional information about the definition. Model-specific indicates that this definition is only applicable to one specific model. The definition of Chesson (2003) is only applicable to models with only one limiting factor (Barabás *et al.*, 2018), as for other models $\mathcal{N}$ and $\mathcal{F}$ depend on the chosen limiting factor. Scaling indicates, that this definition

| Source | Definition | Range | Pos. Eff. | Comp. | Coex. | Add. |
|---|---|---|---|---|---|---|
| Chesson 2003 | $\mathcal{N} = \overline{\left(\frac{\Delta I - \Delta N}{d}\right)}$<br>$\mathcal{F}_i = \frac{r_i}{d_i} - \mathcal{N}$ | $[-\infty, \infty]$<br>$[-\infty, \infty]$ | $[-\infty, \infty]$ | $[-\infty, \infty]$ | $-\mathcal{F}_i \leq \mathcal{N}$ | 1 lim. factor |
| Carroll et al. 2011 | $\mathcal{N}$ =1-Mean(sens.)<br>$\mathcal{F}$ =Std(sens.) | $[-\infty, 1]$<br>$[1, \infty]$ | Undef. | $[-\infty, 1]$ | $\mathcal{F} \leq \frac{1}{1-\mathcal{N}}$ | Multi-species |
| Zhao et al. 2016 | $\mathcal{N} = 1 + r_i + r_j$<br>$\mathcal{F}_i = \log_{10}\left(\frac{K_i}{K_j}\right)$ | $[-\infty, \infty]$<br>$[-\infty, \infty]$ | $[-\infty, \infty]$ | $[-\infty, \infty]$ | None | Scaling |
| Carmel et al. 2017 | $\frac{2-\mathcal{N}}{2\sqrt{1-\mathcal{N}}} = \text{Mean}\left(\frac{r_i}{\mu_i} + 1\right)$<br>$\mathcal{F} = \left(\text{Std}\left(\frac{r_i}{\mu_i} + 1\right)\right)^2$ | $[0, 1]$<br>$[1, \infty]$ | [0,1] | Undef. | $\mathcal{F} \leq \left(\frac{2-\mathcal{N}}{2\sqrt{1-\mathcal{N}}}\right)^2$ | Multi-species |
| Saavedra et al. 2017 | $\mathcal{N} = \frac{2}{\pi}\arcsin\left(\frac{\alpha_{ii}\alpha_{jj} - \alpha_{ij}\alpha_{ji}}{\sqrt{\alpha_{ii}^2 + \alpha_{ji}^2}\sqrt{\alpha_{jj}^2 + \alpha_{ij}^2}}\right)$<br>$\mathcal{F} = \frac{180}{\pi}\arccos\left(\frac{r \cdot r_c}{\lVert r \rVert \cdot \lVert r_c \rVert}\right)$ | $[0, 1]$<br>$[0, 90]$ | Undef<br>$[0, 1]$ | $\mathcal{N} < 0$ | $\mathcal{F} \leq 45 \cdot \mathcal{N}$ | Scaling<br>Multi-species |
| Godoy & Levine 2014 | $\mathcal{N} = 1 - \sqrt{\frac{a_{ij}a_{ji}}{a_{ii}a_{jj}}}$<br>$\mathcal{F}_i = \frac{\lambda_i - 1}{\lambda_j - 1}\sqrt{\frac{a_{ji}a_{jj}}{a_{ij}a_{ii}}}$ | $[0, 1]$<br>$[0, \infty]$ | Undef. | $\mathcal{N} < 0$ | $\mathcal{F}_i \leq \frac{1}{1-\mathcal{N}}$ | Model-specific |
| Adler et al. 2007 | $\mathcal{N}_i = \log\left(\frac{\lambda_j}{1 + \frac{a_{ij}}{a_{jj}}(\lambda_j - 1)}\right)$<br>$\mathcal{F}_i = \log\left(\frac{\lambda_i}{\lambda_j}\right)$ | $[0, \infty]$<br>$[-\infty, \infty]$ | $[0, \infty]$ | $\mathcal{N} < 0$ | $-\mathcal{F}_i \leq \mathcal{N}_i$ | Model-specific |
| Bimler et al. 2018 | $\mathcal{N} = 1 - \frac{e^{a_{ij} + a_{ji}}}{e^{a_{ii} + a_{jj}}}$<br>$\mathcal{F}_i = \frac{e^{a_{ji} + a_{jj}}}{e^{a_{ii} + a_{ij}}}$ | $[0, 1]$<br>$[0, \infty]$ | $[0, 1]$ | $\mathcal{N} < 0$ | None | Model-specific, Scaling |
| Chesson 1990 | $\mathcal{N} = 1 - \sqrt{\frac{a_{ij}a_{ji}}{a_{ii}a_{jj}}}$<br>$\mathcal{F}_i = \sqrt{\frac{a_{ji}a_{jj}}{a_{ij}a_{ii}}}$ | $[0, 1]$<br>$[0, \infty]$ | Undef. | $\mathcal{N} < 0$ | $\mathcal{F}_i \leq \frac{1}{1-\mathcal{N}}$ | Model-specific |
| Chesson & Kuang 2008 | $\mathcal{N} = 1 - \frac{\sqrt{\alpha_{ij}^R\alpha_{ji}^R} + \sqrt{\alpha_{ij}^P\alpha_{ji}^P}}{s_i s_j}$<br>$\mathcal{F}_i = \frac{s_j}{s_i}\frac{\mu_i^R - \mu_i^P - m_i}{\mu_j^R - \mu_j^P - m_j}$ | $[0, 1]$<br>$[0, \infty]$ | Undef. | $\mathcal{N} < 0$ | $\mathcal{F}_i \leq \frac{1}{1-\mathcal{N}}$ | Model-specific |
| Spaak & DeLaender | $\mathcal{N}_i = \frac{f_i(0, N_j^*) - f_i(N_j^*, 0)}{f_i(0,0) - f_i(c_j N_j^*, 0)}$<br>$\mathcal{F}_i = \frac{f_i(N_j^*, 0)}{f_i(0,0)}$ | $[0, 1]$<br>$[-\infty, 1]$ | $\mathcal{N} > 1$ | $\mathcal{N} < 0$ | $\mathcal{F}_i \leq \frac{\mathcal{N}_i}{1-\mathcal{N}_i}$ | - |

**Table A1:** Summary of the different definitions of $\mathcal{F}$ and $\mathcal{N}$ in the literature. Red entries denote undesirable behaviour. For further explanation see text.

depends on the parametrisation of the model. The easiest way to see this is Zhao *et al.* (2016). If the same experiment were performed twice, once measured in $mg$ and once in $\mu l$, the $\mathcal{F}$ would differ, as the two bacteria strains will not have the same physical density ($\frac{mg}{\mu l}$). Similarly in Bimler *et al.* (2018) the values of $\mathcal{N}$ and $\mathcal{F}$ change if we re-parametrise the system to have $\alpha_{ii} = 1$. The structural equivalent of fitness differences in Saavedra *et al.* (2017) is also affected if we re-parametrise to have $r_i = 1$. Finally the definitions of Carmel *et al.* (2017); Carroll *et al.* (2011) and Saavedra *et al.* (2017) can be applied to multispecies communities, however in this case the coexistence condition does not suffice to assess coexistence of the species in a community. That is there might be two communities, that have the same $\mathcal{N}$ and $\mathcal{F}$ values, but the species of one community will coexist, while the species in the other will not.

#### Appendix B: Mathematical proofs

Before we go into the proofs and special cases we first introduce some appendix specific notation. As in the main text we use the following notation to describe population dynamics:

$$\frac{1}{N_i}\frac{dN_i}{dt} = f_i(N_i, N_j) \tag{B1}$$

Note that this notation is slightly different from the usual form, which is

$$\frac{1}{N_i}\frac{dN_i}{dt} = f_i(N_1, N_2) \tag{B2}$$

In our notation the density of the focal species is the first argument of the growth function of said species. Opposed to the more conventional natural order of the species, this allows us to more easily write $f_i(0, N_j)$ and similar expressions. The growth rates are assumed to be biological, explicitly we assume

$$1. f_i \text{ are continuous functions in all their arguments} \tag{B3}$$

$$2. \exists \lim_{N_i \to 0} f_i(N_i, 0) = f_i(0,0) \tag{B4}$$

$$3. \limsup_{N_i \to \infty} f_i(N_i, N_j) < 0 \tag{B5}$$

The first assumption is obviously fulfilled for any biological system. The second assumption states that it is reasonable to talk about the monoculture growth rate $f_i(0,0)$. The third assumption ensures that species cannot reach unlimited densities. Those not familiar with the limes supremum can just replace the limes supremum with the normal limes. The limit is taken with fixed but arbitrary $N_j$. Especially we do not assume that the limit is equal for different $N_j$ or that the limit has to be $-\infty$, nor is anything said about the uniformity of the convergence.

We add the following assumption

$$4. f_i(0,0) > f_i(N_i, 0) \tag{B6}$$

That is the per capita growth rate in monoculture is maximal at minimal density. While most biological systems fulfill this assumption it is not a biological necessity. It is solely included for mathematical purposes.

Finally we introduce the following notation

$$\mathcal{N}_1(c) = \frac{f_1(0, N_2^*) - f_1(cN_2^*, 0)}{f_1(0,0) - f_1(cN_2^*, 0)} \tag{B7}$$

$$\mathcal{N}_2(c) = \frac{f_2(0, N_1^*) - f_2(\frac{1}{c}N_1^*, 0)}{f_2(0,0) - f_2(\frac{1}{c}N_1^*, 0)} \tag{B8}$$

Where the niche difference depends on the conversion factor $c = c_2$. We omitted the dependence of $c$ on the species, as we have seen that $c_i = \frac{1}{c_j}$. This definition is asymmetric in the species, however the arbitrary choice that species 1 depends linearly on $c$ only affects $c$ itself, not the $\mathcal{N}$.

#### *Existence of $\mathcal{N}$ and c*

We want to solve $|1-\mathcal{N}_i(c)| = |1-\mathcal{N}_j(c)|$ for $c$, we therefore have to prove that such a solution always exists. The solution will be denoted $c'$. In this part we only prove the existence of $c'$, the uniqueness of such a $c'$ will be discussed below. We first present a "proof" based on biological meaning and logic, that requires a minimum of mathematical knowledge. In this "proof" we omit the absolute values. This biological reasoning is then turned into a rigorous mathematical proof.

The definition of $\mathcal{N}$ essentially compares the interspecific competition ($f_1(0, N_2^*)$) with intraspecific competition ($f_1(cN_2^*, 0)$). The intraspecific competition however depends on the conversion factor $c$, which can take any positive value. The larger this $c$ is, the larger the intraspecific competition will be. By choosing the $c$ being 0 (respectively $\infty$) intraspecific competition will be very small (large) compared to interspecific competition and hence $\mathcal{N}$ will be $-\infty$ ($> 0$). The species however react differently to this value $c$, such that we have a value $c''$ where $\mathcal{N}_1 < \mathcal{N}_2$ and a value $c'''$ with $\mathcal{N}_2 < \mathcal{N}_1$, therefore there must also be a value, where they are equal.

**Theorem 1.** *$f_i(0,0) \neq f_i(0, N_j^*) \Rightarrow \exists c' : |1-\mathcal{N}_1(c')| = |1-\mathcal{N}_2(c')|$*

*Proof.* We define $A_i = \lim_{N\to\infty} f_i(N, 0)$. For simplicity we assume the existence of the limit, if this is not given a similar proof can be done with lim sup. With this definition we can compute:

$$\lim_{x\to\infty} |1-\mathcal{N}_1(x)| = \left|\lim_{x\to\infty} \frac{f_1(0,0) - f_1(0,N_2^*)}{f_1(0,0) - f_1(xN_2^*,0)}\right| \tag{B9}$$

$$= \left|\frac{f_1(0,0) - f_1(0,N_2^*)}{f_1(0,0) - \lim_{x\to\infty} f_1(xN_2^*,0)}\right| \tag{B10}$$

$$= \left|\frac{f_1(0,0) - f_1(0,N_2^*)}{f_1(0,0) - A_1}\right| \tag{B11}$$

$$< \infty \tag{B12}$$

$$\lim_{x\to 0}|1-\mathcal{N}_1(x)| = \left|\lim_{x\to 0}\frac{f_1(0,0)-f_1(0,N_2^*)}{f_1(0,0)-f_1(xN_2^*,0)}\right| \tag{B13}$$

$$= |f_1(0,0)-f_1(0,N_2^*)| \lim_{y\to f_1(0,0)} \frac{1}{|f_1(0,0)-y|} \tag{B14}$$

$$= \infty \tag{B15}$$

With similar arguments we get $\lim_{c\to 0} 1-\mathcal{N}_2(c) < \infty, \lim_{c\to\infty}\mathcal{N}_2(c) = \infty$. By the intermediate value theorem we therefore have the existence of a $c'$ with equality. □

Now to the special case $f_i(0,0) = f_i(0,N_j^*)$. This implies $\mathcal{N}_i = 1$ independent of our choice of $c$. If it happens to be that we also have $f_j(0,0) = f_j(0,N_i^*)$ we also have $\mathcal{N}_j = 1$ for all $c$, which solves the problem. However in this case we can't compute the $c$ and hence not compute the $\mathcal{F}$. We therefore define $\mathcal{F} = 1$ for both species in this case.

In most of the cases however we will have $f_j(0,0) \neq f_j(0,N_j^*)$. The equation to be solved then becomes $\left|\frac{f_j(0,0)-f_j(0,N_i^*)}{f_j(0,0)-f_j(cN_i^*,0)}\right| = 0$, which has the "solution" $f_j(cN_i^*,0) = -\infty, c = \infty$. All we therefore have to allow is setting $c = \infty$. With this we can compute $\mathcal{F}_i = \frac{f_i(\frac{1}{\infty}N_j^*,0)}{f_i(0,0)} = 1$ and $\mathcal{F}_j = \frac{f_j(\infty,0)}{f_j(0,0)} = -\infty$. We implicitly assume that $\lim_{N\to\infty} f_i(N,0) = -\infty$, which seems to be a biologically reasonable assumption. This is equivalent to previous definitions (Carroll *et al.*, 2011; Chesson, 2000; Godoy & Levine, 2014). Note that the coexistence condition $-\mathcal{F}_j \leq \frac{\mathcal{N}_j}{1-\mathcal{N}_j}$ becomes useless, as both sides are $\infty$.

#### *Uniqueness of $\mathcal{N}$ and c*

In general the equation $|1-\mathcal{N}_i(c)| = |1-\mathcal{N}_j(c)|$ might not have a unique solution $c'$ but rather multiple solutions $c'_k$. The equation $|1-\mathcal{N}_i(c)| = |1-\mathcal{N}_j(c)|$ can be arbitrarily complicated, such that in general we can't tell in advance whether there is a unique solution. However we can give a sufficient condition under which the solution is unique, this condition also happens to be biologically meaningful: If both species have strictly negative density dependencies in monoculture (which most communities have), then the $c'$ and also the $\mathcal{N}$ will be unique. Essentially the monotonicity of the monoculture per capita growth rates $\frac{\partial f_i(N_i,0)}{\partial N_i} < 0$ translates into a monotonicity of

$1 - \mathcal{N}_i$, by this monotonicity there can be only one $c'$ with equality.

**Theorem 2.** *If we assume that* $\frac{\partial f_i(N_i,0)}{\partial N_i} < 0$ *then the definition of* $\mathcal{N}_i$ *and of* $c'$ *is unique.*

*Proof.* We simply have to compute the derivative:

$$\frac{d|1-\mathcal{N}_1(x)|}{dx} = \frac{d}{dx}\left(\frac{|f_1(0,0) - f_1(0,N_2^*)|}{|f_1(0,0) - f_1(xN_2^*,0)|}\right) \tag{B16}$$

$$= |f_1(0,0) - f_1(0,N_2^*)| \cdot (-1) \tag{B17}$$

$$\cdot \left(f_1(0,0) - f_1(xN_2^*,0)\right)^{-2} \cdot (-1)\frac{\partial f_1}{\partial N_1} \cdot N_2^* \tag{B18}$$

$$= N_2^* \frac{|f_1(0,0) - f_1(0,N_2^*)|}{(f_1(0,0) - f_1(xN_2^*,0))^2}\frac{\partial f_1}{\partial N_1} < 0 \tag{B19}$$

We removed the absolute values from $f_i(0,0) - f_i(xN_2,0)$, as this is always positive according to assumption 4. With similar computation we see that $\frac{d|1-\mathcal{N}_2(x)|}{dx} > 0$ and therefore there exists only one $c'$ □

#### *Multispecies case*

The ideas used in the multispecies case are essentially equivalent to the two species case. The growth model is now assumed to have the following shape:

$$\frac{1}{N_i}\frac{dN_i}{dt} = f_i(N_i, \mathbf{N}^{-\mathbf{i}}) \tag{B20}$$

Where $\mathbf{N}^{-\mathbf{i}}$ is the vector of all species densities with the density of species $i$ removed. The second argument of $f_i$ will in this part be a vector of dimension $n-1$ and be **bold**. We then define

$$\mathcal{N}_i = \frac{f_i(0, \mathbf{N}^{-\mathbf{i},*}) - f_i(\sum_{j\neq i} c_{ij}N_j^{-i,*}, \mathbf{0})}{f_i(0,\mathbf{0}) - f_i(\sum_{j\neq i} c_{ij}N_j^{-i,*}, \mathbf{0})} \tag{B21}$$

$$\mathcal{F}_i = \frac{f_i(\sum_{j\neq i} c_{ij}N_j^{-i,*}, \mathbf{0})}{f_i(0,\mathbf{0})} \tag{B22}$$

with $\mathbf{0}$ is the $n-1$ dimensional 0 vector and $N_j^{-i,*}$ is the equilibrium density of species $j$ (in the presences of the $n-1$ resident species) and $c_{ij}$ is the conversion factor from species $j$ to species $i$.

The $c_{ij}$ can be found with the two species communities. That is for each pair of species we compute the $\mathcal{N}$ of those two species in the presence of all the other species denoted $\mathcal{N}_{ij}$. The presence of the other species is important to include the indirect and higher order effects.

$$\mathcal{N}_{ij} = \frac{f_i(0, \mathbf{N^{-i,*}}) - f_i(c_{ij}N_j^{-i,*}, \mathbf{N^{-i,*}_{j\to 0}})}{f_i(0, \mathbf{N^{-i,*}_{j\to 0}}) - f_i(c_{ij}N_j^{-i,*}, \mathbf{N^{-i,*}_{j\to 0}})} \tag{B23}$$

Where $N_{j\to 0}^{-i,*}$ is the equilibrium density of the community in the absence of species $i$ with the density of species $j$ set to zero. This is not to be confused with $N^{-ij,*}$ the equilibrium density of the community in the absence of species $i$ and $j$. That is to obtain $N_{j\to 0}^{-i,*}$ we first compute $N^{-i,*}$ and then set density of species $j$ to zero, without affecting the densities of the other species. Notationally the $*$ is not optimal, as this is not an equilibrium of the system. Then we solve all the equations $|1 - \mathcal{N}_{ij}| = |1 - \mathcal{N}_{ji}|$ to obtain $c_{ij}$, which allows us to compute the total $\mathcal{N}_i$. The same proofs for existence and uniqueness apply for the multispecies case.

#### *Decomposition of invasion growthrate*

$$\mathcal{N}_i + \mathcal{F}_i - \mathcal{N}_i \cdot \mathcal{F}_i = \frac{f_i(0, N_j^*) - f_i(c_jN_j^*, 0)}{f_i(0,0) - f_i(c_jN_j^*, 0)} + \frac{f_i(c_jN_j^*, 0)}{f_i(0,0)} \tag{B24}$$

$$- \frac{f_i(0, N_j^*) - f_i(c_jN_j^*, 0)}{f_i(0,0) - f_i(c_jN_j^*, 0)} \cdot \frac{f_i(c_jN_j^*, 0)}{f_i(0,0)} \tag{B25}$$

$$= \frac{f_i(0, N_j^*)f_i(0,0) - f_i(c_jN_j^*, 0)f_i(0,0)}{(f_i(0,0) - f_i(c_jN_j^*, 0))f_i(0,0)} \tag{B26}$$

$$+ \frac{f_i(c_jN_j^*, 0)f_i(0,0) - f_i(c_jN_j^*, 0)^2}{(f_i(0,0) - f_i(c_jN_j^*, 0))f_i(0,0)} \tag{B27}$$

$$- \frac{f_i(0, N_j^*)f_i(c_jN_j^*, 0) - f_i(c_jN_j^*, 0)^2}{(f_i(0,0) - f_i(c_jN_j^*, 0))f_i(0,0)} \tag{B28}$$

$$= \frac{f_i(0, N_j^*)f_i(0,0) - f_i(0, N_j^*)f_i(c_jN_j^*, 0)}{(f_i(0,0) - f_i(c_jN_j^*, 0))f_i(0,0)} \tag{B29}$$

$$= \frac{f_i(0, N_j^*)}{f_i(0,0)} \tag{B30}$$

### Appendix C: Examples

#### *Mac-Arthur resource model*

First we deduce the Lotka-Volterra model from the resource model of MacArthur (1970).

As stated in the main text, the growth rates for the consumer species and the resources respectively are:

$$\frac{1}{N_i}\frac{dN_i}{dt} = \sum_{l=1}^{m} u_{il}R_l - m_i \tag{C1}$$

$$\frac{1}{R_l}\frac{dR_l}{dt} = K_l - R_l - \sum_{i=1}^{n} u_{il}N_i \tag{C2}$$

Where $N_i$ is the consumer density, $u_{il}$ is the rate at which species $i$ consumes resource $l$, $R_l$ is the density of resource $l$, and $m_i$ is the loss rate and $K_l$ is the carrying capacity.

We assume that the dynamics of $R_l$ are much faster than the dynamics of $N_i$, that is we explicitly assume that $R_l$ is always at equilibrium, i.e. $R_l = K_l - \sum_{j=1}^{n} u_{jl}N_j$. Inserting this into the species growth rates we get:

$$\begin{aligned}
\frac{1}{N_i}\frac{dN_i}{dt} &= \sum_{l=1}^{m} u_{il}\left(K_l - \sum_{j=1}^{n} u_{jl}N_j\right) - m_i && \text{(C3)}\\
&= \sum_{l=1}^{m} u_{il}K_l - m_i - \sum_{l=1}^{m}\sum_{j=1}^{n} u_{jl}u_{il}N_j && \text{(C4)}\\
&= \underbrace{\sum_{l=1}^{m} u_{il}K_l - m_i}_{\mu_i} - \sum_{j=1}^{n} N_j \underbrace{\sum_{l=1}^{m} u_{il}u_{jl}}_{\langle u_i, u_j\rangle} && \text{(C5)}\\
&= \mu_i - \sum_{j=1}^{n}\langle u_i, u_j\rangle N_j && \text{(C6)}
\end{aligned}$$

Usually the species specific interaction $\langle u_i, u_j\rangle$ is denoted $\alpha_{ij}$ in the Lotka-Volterra model, however, we choose the scalar product notation because it gives a clear biological interpretation of $\mathcal{N}$, $\mathcal{F}$ and $c$. We continue to compute the $\mathcal{N}$ and $\mathcal{F}$ for the *two* species case (for multispecies case see Appendix C, Multispecies): The monoculture equilibria are $N_i^* = \frac{\mu_i}{\alpha_{ii}}$ and the $\mathcal{N}(c)$ (using

$f_i(N_i, N_j) = \mu_i - \sum_{j=1}^n \langle u_i, u_j \rangle N_j)$:

$$\mathcal{N}_1(c) = \frac{(\mu_1 - \frac{\langle u_1, u_2 \rangle}{\langle u_2, u_2 \rangle}\mu_2) - (\mu_1 - c\frac{\langle u_1, u_1 \rangle}{\langle u_2, u_2 \rangle}\mu_2)}{\mu_1 - (\mu_1 - c\frac{\langle u_1, u_1 \rangle}{\langle u_2, u_2 \rangle}\mu_2)} \tag{C7}$$

$$= \frac{-\frac{\langle u_1, u_2 \rangle}{\langle u_2, u_2 \rangle}\mu_2 + c\frac{\langle u_1, u_1 \rangle}{\langle u_2, u_2 \rangle}\mu_2}{c\frac{\langle u_1, u_1 \rangle}{\langle u_2, u_2 \rangle}\mu_2} = 1 - \frac{1}{c}\frac{\langle u_1, u_2 \rangle}{\langle u_1, u_1 \rangle} \tag{C8}$$

$$\mathcal{N}_2(c) = 1 - c\frac{\langle u_2, u_1 \rangle}{\langle u_2, u_2 \rangle} \tag{C9}$$

Solving for $c$ yields $c = \frac{\|u_2\|}{\|u_1\|}$, remember that $c = c_2$, i.e. we have $c_i = \frac{\|u_i\|}{\|u_j\|}$. Which in turn gives $\mathcal{N}_1 = \mathcal{N}_2 = 1 - \frac{\langle u_1, u_2 \rangle}{\|u_1\| \cdot \|u_2\|}$, where $\|u_i\| = \sqrt{\langle u_i, u_i \rangle}$. Note that this definition is equivalent to Chesson (1990) but a completely independent proof with a different interpretation. The fitness differences are $\mathcal{F}_i = 1 - \frac{\mu_j}{\mu_i}\frac{\|u_i\|}{\|u_j\|}$.

#### *General resource uptake model*

In the previous section we showed that $c_i = \frac{\|u_i\|}{\|u_j\|}$ scales the total resource consumption of the two species in the MacArthur resource model. In this section we assume a more general version of a resource specific model and show that $c_i$ scales the resource uptake of both species. We assume the following growth rates for the species and the resources:

$$\frac{1}{N_i}\frac{dN_i}{dt} = g_i(u_i(R)) - m_i \tag{C10}$$

$$\frac{dR_l}{dt} = G_l(R) - \sum_i u_i^l(R)N_i \tag{C11}$$

Where $u_i^l(R)$ is the per capita consumption of resource $l$ by the species $i$. $u_i(R) = (u_i^l(R))_l$ is the vector containing the consumption of all resources. $g_i$ is the conversion of resources eaten by the species into biomass, $m_i$ is the mortality rate of the species and $G_l$ is the regeneration function of the resource $R_l$, which may depend on the densities of the other resources. We assume that the only interaction between species and resources is consumption and species interact only indirectly with each other via depletion of resources. Furthermore we assume that the dynamics of the resources are much faster and the density of the resources are always in equilibrium, denoted $R^*(N)$. Finally we assume that the functions $u_i, g_i$ and $R^*$ are monotone, more specifically, the more resources there are, the more the species consumes, the more it consumes the faster it growth (i.e. $\frac{\partial g_i}{\partial R_l} \geq 0$ and $\frac{\partial u_i^l}{\partial R_{l'}} \geq 0$). On the other hand the higher the species densities the lower the resource levels (i.e. $\frac{\partial R_l^*}{\partial N_{i,j}} \leq 0$).

As we do not take any more specific assumptions on the functions $g_i, u_i$ and $G_l$ we can't compute $c_i$ explicitly to proof that $c_i$ scales the total amount of resources consumed. Instead we ask whether $c_i$ increases when species $i$ consumes more. Implicitly the resource uptake functions $u_i$ depends on traits $t_j^i$. We focus on one specific trait $t_1$ and assume that higher values of this trait imply higher consumption (i.e. $\frac{\partial u_i^l}{\partial t_1} > 0$). We show that a species $i'$ which consumes more (i.e. $t_1' > t_1$) will have a lower $c_i$ (i.e. $\frac{\partial c_i}{\partial t_1} < 0$) using the implicit function theorem. The parameter $c_i$ therefore becomes a function of $t_1$.

$c_i^0$ and $t_1^0$, the original values for $c_i$ and the trait $t_1$, are a solution to the following equation

$$1 - \frac{f_i(0,0) - f_i(0, N_j^*)}{f_i(0,0) - f_i\left(\frac{N_j^*}{c_i}, 0\right)} = 1 - \frac{f_j(0,0) - f_j(0, N_i^*)}{f_i(0,0) - f_i(c_i N_i^*, 0)} \tag{C12}$$

$$\underbrace{\frac{g_i(u_i(t_1, K)) - g_i(u_i(t_1, R_{t_1}^*(0, N_j^*)))}{g_i(u_i(t_1, K)) - g_i(u_i(t_1, R_{t_1}^*(c_i^{-1} N_j^*, 0)))}}_{h_i(c_i, t_1)} = \underbrace{\frac{g_j(u_j(K)) - g_j(u_j(R_{t_1}^*(0, N_i^*(t_1))))}{g_j(u_j(K)) - g_j(u_j(R_{t_1}^*(c_i N_i^*(t_1), 0)))}}_{h_j(c_i, t_1)} \tag{C13}$$

Hence we define $h(c_i, t_1) = h_i(c_i, t_1) - h_j(c_i, t_1)$. We know that $h\left(c_i^0, t_1^0\right) = 0$ and we can therefore use the implicit function theorem to compute $\frac{\partial c_i}{\partial t_1} = -\left(\frac{\partial h}{\partial c_i}\right)^{-1} \frac{\partial f}{\partial t_1}$.

$$\frac{\partial h_i}{\partial c_i} = -\frac{g_i(u_i(t_1, K)) - g_i(u_i(t_1, R_{t_1}^*(0, N_j^*)))}{\left(g_i(u_i(t_1, K)) - g_i(u_i(t_1, R_{t_1}^*(c_i^{-1} N_j^*)))\right)^2} \tag{C14}$$

$$\cdot \frac{\partial}{\partial c_i}\left(g_i(u_i(t_1, R_{t_1}^*(c_i^{-1} N_j^*, 0)))\right) \tag{C15}$$

The first factor is always positive as $K > R_{t_1}^*(0, N_j^*)$. The second factor is positive, as increasing $c_i$ decreases species abundances ($c_i^{-1} N_j^*$), which increases resource abundance ($R_{t_1}^*$), which increases resource consumption ($u_i$), which increases biomass accumulation($g_i$), hence $\frac{\partial h_i}{\partial c_i} < 0$. Similar arguments show that $\frac{\partial h_j}{\partial c_i} > 0$ and hence $\frac{\partial h}{\partial c_i} = \frac{\partial h_i}{\partial c_i} - \frac{\partial h_j}{\partial c_i} < 0$.

To compute $\frac{\partial h_i}{\partial t_1}$ we take the following assumption

$$\frac{\partial}{\partial t_1}\left(g_i(u_i(t_1, R^*_{t_1}(0, N_j^*)))\right) > \frac{\partial}{\partial t_1}\left(g_i(u_i(t_1, R^*_{t_1}(c_i^{-1}N_j^*, 0)))\right) \tag{C16}$$

In the left hand side increasing $t_1$ will increase the consumption and therefore the conversion to biomass. However on the right hand side increasing $t_1$ will decrease the resource levels, as species $i$ will consume more. This leads to $\frac{\partial h_i}{\partial t_1} < 1$. Similarly we can show that $\frac{\partial h_j}{\partial t_1} > 1$ which leads to $\frac{\partial h}{\partial t_1} < 0$. As a consequence we have $\frac{\partial c}{\partial t_1} = -\left(\frac{\partial h}{\partial c}\right)^{-1} \cdot \frac{\partial h}{\partial t_1} < 0$, i.e. under the assumptions taken, higher consumption of resources leads to lower $c_i$ and therefore $c_i$ can be seen as a scaling factor between the total amount of resources consumed of the two species.

#### *Positive interspecific interactions*

Generating a resource implies that a species has a negative utilisation of that resource (similar for limiting predator efficacy), which would correspond to a negative bar in figure **??**. If the total effect of species $j$ on species $i$ is positive (facilitation), then species $j$ positively affects the limiting factors. $c_j$ is a measure of how *much* species $j$ affects the environment, independent of whether such an effect is positive or negative. We can therefore obtain $c_j$ by making the absolute values of the effects on the limiting factors equal, i.e. $|1-\mathcal{N}_i| = |1-\mathcal{N}_j|$.

To investigate the case where one species facilitates the other we tweak the resource model slightly by allowing species 1 to generate a resource $P$ that can be used by species 2 for its growth. The differential equations become

$$\frac{1}{N_1}\frac{dN_1}{dt} = \left(\sum_{l=1}^{m} u_{1l}R_l - m_1 - p_1\right) \tag{C17}$$

$$\frac{1}{N_2}\frac{dN_2}{dt} = \left(\sum_{l=1}^{m} u_{2l}R_l - m_2 + u_{2p}P\right) \tag{C18}$$

$$\frac{1}{R_l}\frac{dR_l}{dt} = K_l - R_l - \sum_{i=1}^{n} u_{il}N_i \tag{C19}$$

$$\frac{dP}{dt} = p_1N_1 - u_{2p}N_2P - kP \tag{C20}$$

The parts in red are different from the usual Mac Arthur model. Species 1 creates resource $P$ with efficiency $p_1$, we from now on however will assume that $p_1$ is incorporated into $m_1$ and omit it. $u_{2p}$ is the utilisation of $P$ by species 2 and $k$ is the decay rate of $P$. We again assume, that the usual resources and the resource $P$ are at equilibrium, which leads to the following equations (for better readability we set $\alpha_{ij} = \langle u_i, u_j \rangle$):

$$\frac{1}{N_1}\frac{dN_1}{dt} = (\mu_1 - \alpha_{11}N_1 - \alpha_{12}N_2) \tag{C21}$$

$$\frac{1}{N_2}\frac{dN_2}{dt} = \left(\mu_2 - \alpha_{21}N_1 - \alpha_{22}N_2 + u_{2p}\frac{p_1 N_1}{k + u_{2p}N_2}\right) \tag{C22}$$

Solving $1 - \mathcal{N}_1 = 1 - \mathcal{N}_2$:

$$1 - \frac{\mu_1 - \left(\mu_1 - \alpha_{12}\frac{\mu_2}{\alpha_{22}}\right)}{\mu_1 - \left(\mu_1 - c\alpha_{11}\frac{\mu_2}{\alpha_{22}}\right)} = 1 - \frac{\mu_2 - \left(\mu_2 - (\alpha_{21} - \frac{u_{2p}p_1}{k})\frac{\mu_1}{\alpha_{11}}\right)}{\mu_2 - \left(\mu_2 - \frac{1}{c}\alpha_{22}\frac{\mu_1}{\alpha_{11}}\right)} \tag{C23}$$

$$\frac{\alpha_{12}\frac{\mu_2}{\alpha_{22}}}{c\alpha_{11}\frac{\mu_2}{\alpha_{22}}} = \frac{\left(\alpha_{21} - \frac{u_{2p}p_1}{k}\right)\frac{\mu_1}{\alpha_{11}}}{\frac{1}{c}\alpha_{22}\frac{\mu_1}{\alpha_{11}}} \tag{C24}$$

$$\frac{\alpha_{12}}{c\alpha_{11}} = c\frac{\alpha_{21} - \frac{u_{2p}p_1}{k}}{\frac{1}{c}\alpha_{22}} \tag{C25}$$

$$c = \sqrt{\frac{\alpha_{22}\alpha_{12}}{\alpha_{11}\left(\alpha_{21} - \frac{u_{2p}p_1}{k}\right)}} \tag{C26}$$

So far this result is only applicable when we assume that $\alpha_{21} - \frac{u_{2p}p_1}{k} \geq 0$, that is we have $f_2(0,0) \geq f_2(0, N_1^*)$ and we are still in the realm of the usual definition. Translating this equation back to the mechanistic model we get $c = \frac{\|u_2\|}{\|u_1\|\sqrt{1 - \frac{u_{2p}p_1}{k\langle u_1, u_2\rangle}}}$, i.e. resource $P$ should indeed be seen as a negative resource and the total amount of resources consumed by species 1 is $\|u_1\|^2 \cdot (1 - \frac{u_{2p}p_1}{k\langle u_1, u_2\rangle})$.

As we have seen in the previous example, the conversion factor are chosen such that both species consume the same total amount of resources. They can however only scale the amount of resources used by a species and not change the sign of the total resources used. In the case $1 - \frac{u_{2p}p_1}{k\langle u_1, u_2\rangle} < 0$ species 1 consumes a negative amount of resources. The conversion factor $c$ can therefore not equate the total amount of resources used by the two species. We therefore equate the absolute value of the total amount of resources used by the two species and therefore also equate $|1-\mathcal{N}_1| = |1-\mathcal{N}_2|$, which leads to $c = \sqrt{\frac{\alpha_{22}\alpha_{12}}{\alpha_{11}\left|\alpha_{21}-\frac{u_2 p p_1}{k}\right|}}$. Note however that we only take the absolute value of $1-\mathcal{N}$ to compute the $c$, we do not change the definition of $\mathcal{N}$ itself. This leads to the fact, that not both species have the same $\mathcal{N}$ as usual. Rather we have $\mathcal{N}_1 = 1 - \sqrt{\frac{\alpha_{12}\left|\alpha_{21}-\frac{u_2 p p_1}{k}\right|}{\alpha_{22}\alpha_{11}}} = 1 - |1-\mathcal{N}_2|$.

#### *Multispecies*

As an example we will solve the multispecies Lotka Voltera model, described by equation C6.

$$\mathcal{N}_{ij} = \frac{\left(\mu_i - \sum_{k\neq i}\alpha_{ik}N_k^{-i,*}\right) - \left(\mu_i - \sum_{k\neq i,j}\alpha_{ik}N_k^{-i,*} - \alpha_{ii}c_{ij}N_j^{-i,*}\right)}{\left(\mu_i - \sum_{k\neq i,j}\alpha_{ik}N_k^{-i,*}\right) - \left(\mu_i - \sum_{k\neq i,j}\alpha_{ik}N_k^{-i,*} - \alpha_{ii}c_{ij}N_j^{-i,*}\right)} \tag{C27}$$

$$= \frac{(\alpha_{ii} - \alpha_{ij}N_j^{-i,*}}{c_{ij}\alpha_{ii}N_j^{-i,*}} \tag{C28}$$

$$= 1 - \frac{\alpha_{ij}}{c_j^i\alpha_{ii}} \tag{C29}$$

Setting the density of species $j$ to zero was done by summing only over the indexes $k \neq i, j$ in the respective function evaluations. Similar to the two species case we have $c_k^i = \frac{1}{c_i^k}$ and hence $c_k^i = \sqrt{\left|\frac{\alpha_{kk}\alpha_{ik}}{\alpha_{ii}\alpha_{ki}}\right|}, \mathcal{N}_{ik} = 1 - \text{sign}(a_{ik})\sqrt{\left|\frac{\alpha_{ik}\alpha_{ki}}{\alpha_{ii}\alpha_{kk}}\right|}$.

$$\mathcal{N}_i = 1 - \frac{\mu_i - \left(\mu_i - \sum_{k\neq i}\alpha_{ik}N_k^{-i,*}\right)}{\mu_i - \left(\mu_i - \sum_{k\neq i}c_k^i\alpha_{ii}N_k^{-i,*}\right)} \tag{C30}$$

$$= 1 - \frac{\sum_{k\neq i}\frac{\alpha_{ik}}{a_{ii}}N_k^{-i,*}}{\sum_{k\neq i}c_k^iN_k^{-i,*}} \tag{C31}$$

$$= 1 - \frac{\sum_{k\neq i}(1-\mathcal{N}_{ik})c_k^iN_k^{-i,*}}{\sum_{k\neq i}c_k^iN_k^{-i,*}} \tag{C32}$$

That is the $1-\mathcal{N}_i$ in multispecies community is a weighted sum of the $1-\mathcal{N}_{ik}$ in two species case.

$$\mathcal{F}_i = \frac{\mu_i - \sum_{k\neq i} c_k^i \alpha_{ii} N_k^{-i^*}}{\mu_i} \tag{C33}$$

$$= 1 - \frac{1}{\mu_i} \sum_{k\neq i} \sqrt{\left|\frac{\alpha_{ik}}{\alpha_{ki}}\right|} \sqrt{\alpha_{ii}\alpha_{kk}} N_k^{-i,*} \tag{C34}$$

$$= 1 - \sum_{k\neq i} \frac{\mu_k}{\mu_i} \sqrt{\left|\frac{\alpha_{ik}\alpha_{ii}}{\alpha_{ki}\alpha_{kk}}\right|} \frac{\alpha_{kk}}{\mu_k} N_k^{-i,*} \tag{C35}$$

$$= 1 - \sum_{k\neq i} (1 - \mathcal{F}_i^k) \frac{N_k^{-i,*}}{N_k^*} \tag{C36}$$

The two species case can be recovered from the multispeices case by noticing that $N_k^{-i,*} = \frac{\mu_k}{\alpha_{kk}}$.

After this simple example we want to mention some possible pitfalls. We saw that $\mathcal{N}_{ik}$ of two species is unaffected of the presence of other species in the Lotka-Volterra model. This is because the species do not change their foraging strategies because of the presence of the other species, i.e. there are no higher order effects (Grilli *et al.*, 2017). This will however in general not be the case. Same holds true for the conversion factors $c_k^i$.

### Appendix D: Material and Methods

Experiments were performed in semi-continuous flow through systems in 6-well plates. Light intensity was set at 1500 lux at the top of the 6 well plates and the walls of the wells were painted black to have a unidirectional light gradient (Huismann & Weissing, 1994). The 6 well plates were mounted on a shaker shaking with 150 rpm to have homogenous culture. The experiment was performed at 20°C. Each well was filled with 5ml brackish mineral medium: $NaCl(8.25gl^{-1})$, $MgCl_2 \cdot 6H_2O(0.66gl^{-1})$, $KCl(0.17gl^{-}1)$, $MgSO_4 \cdot 7H_2O$ $(1.16gl^{-1})$, $CaCl_2 \cdot 2H_2O(0.17gl^{-1})$, $Na_3$-citrate $(4.98mgl^{-1})$, $Na_2$-EDTA $(0.83mgl^{-1})$, $NaNO_3(1.25gl^{-1})$, $Na_2CO_3$ $(46.0mgl^{-1})$, trace metal mix $(1.0mgl^{-1})$, $K_2HPO_4 \cdot 3H_2O(33.2mgl^{-1})$, $Fe\text{-}NH_4$-citrate $(4.8mgl^{-1})$ (Stomp *et al.*, 2004). Twice a week (all 84 hours) we added $200\mu l$ distilled water to counteract evaporation and replaced 1ml (20%) of brackish medium. Densities were measured with a flow-cytometer, to distinguish the cells we used an EM-clustering algorithm on the yellow and red fluorescence channels. To inoculate with above equilibrium densities we centrifuged 50ml of culture at 2000g for 20 minutes. Density measures, code to produce the figure and compute the $\mathcal{N}$ and $\mathcal{F}$ values can be found on https://github.com/juergspaak/NFD_definitions.

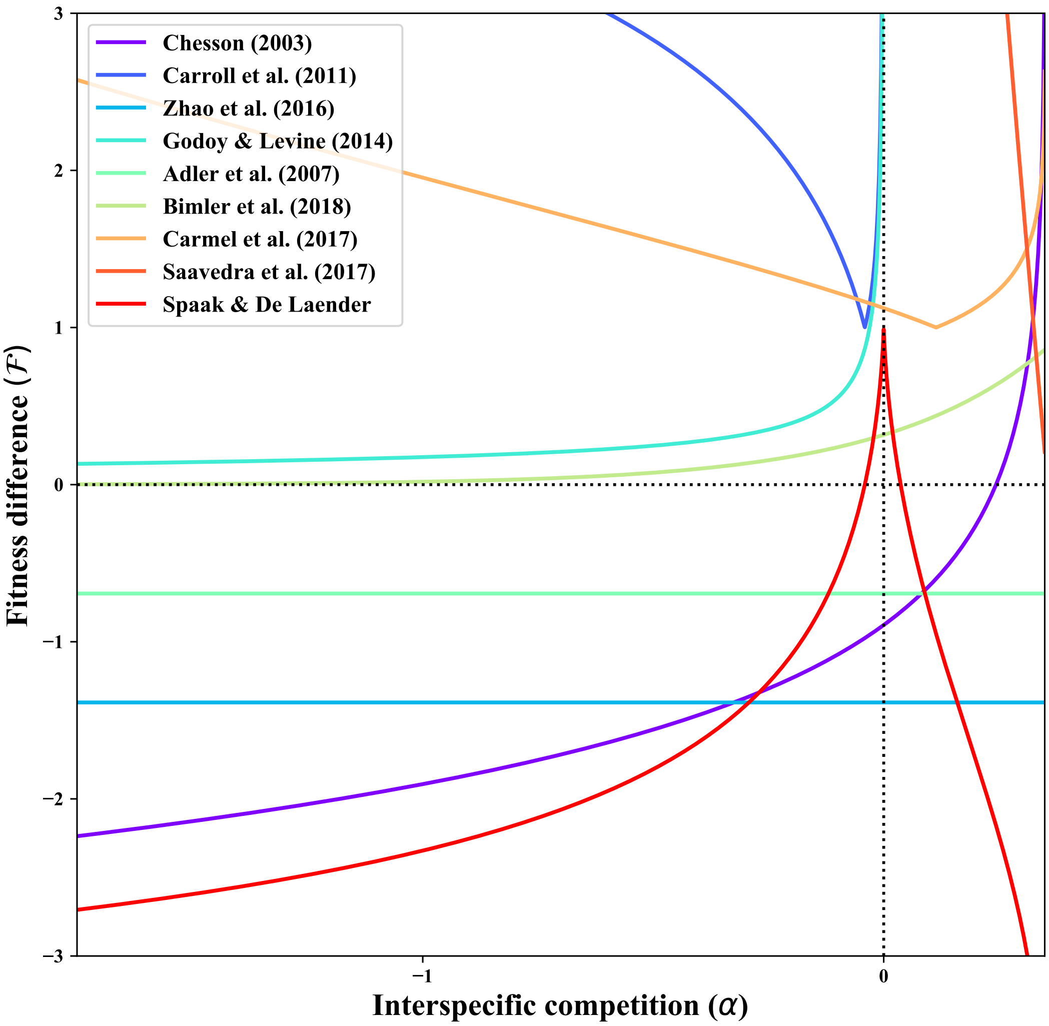


**Figure A 1:** $\mathcal{F}$ values for the annual plant model according to different definitions for varying interspecific competition $\alpha = -\frac{\alpha_{12}\alpha_{21}}{\alpha_{11}\alpha_{22}}$. Comparing the different definitions of $\mathcal{F}$ is more difficult than comparing the different $\mathcal{N}$, as some definitions interpret $\mathcal{F} = 0$ to be equal fitness while other have $\mathcal{F} = 1$ for equal fitness. For the definition of Chesson (2003) we chose species 1 to be the limiting factor. Parameter values are: $\lambda_1 = 1.5, \lambda_2 = 3, \alpha_{11} = 1, \alpha_{22} = 1, \alpha_{21} = 0.7$ and $\alpha_{12}$ varies in $[-0.5, 2.5]$.
